# Members of the *Legionella pneumophila* Sde Family Target Tyrosine Residues for Phosphoribosyl-Linked Ubiquitination

**DOI:** 10.1101/2021.05.05.442833

**Authors:** Mengyun Zhang, Joseph M. McEwen, Nicole M. Sjoblom, Kristin M. Kotewicz, Ralph R. Isberg, Rebecca A. Scheck

## Abstract

*Legionella pneumophila* establishes a replication vacuole by translocating hundreds of protein effectors through a type IV secretion system (T4SS). Among these translocated effectors are members of the Sde family, which catalyze phosphoribosyl-linked ubiquitination (pR-Ub) of host targets. Previous work has posited that Sde proteins solely target serine (Ser) residues within acceptor protein substrates. We show here that SdeC-mediated pR-Ub modification results from a stepwise reaction that also modifies tyrosine (Tyr) residues. Unexpectedly, the presence of an HA tag on Ub resulted in poly-pR-ubiquitination, consistent with the HA tag acting as an acceptor target. Interrogation of phosphoribosyl-linked HA-Ub revealed that Tyr4 was the preferred targeted residue, based on LC-MS/MS analysis of the crosslinked product. Further analysis using synthetic HA variants revealed promiscuous modification of Tyr, as crosslinking was prevented only by constructing a triple mutant in which all three Tyr within the HA sequence were substituted with Phe. Although previous work has indicated that Ser is the sole acceptor residue, we found no evidence of Ser preference over Tyr using Tyr->Ser replacement mutants. This work demonstrates that pR-ubiquitination by the Sde family is not limited to Ser-modification as previously proposed, and broadens the potential sites targeted by this family.

## Introduction

*Legionella pneumophila* is a Gram-negative bacterium that grows intracellularly within a host cell-derived membrane-bound compartment.^1,2^ Intracellular growth and biogenesis of the replication vacuole requires the assembly of the Icm/Dot complex,^3–5^ a type IV secretion system (T4SS) that spans the bacterial envelope and allows translocation of over 300 proteins (called *effectors*, throughout) across the vacuole.^6–10^ The biochemical activities of dozens of these substrates are known to regulate host cell vesicle trafficking,^11–13^ influence tubular endoplasmic reticulum (ER) dynamics,^14–17^ rewire host ubiquitination (Ub),^18,19^ as well inhibit host protein synthesis.^20–22^ Manipulation of these processes largely results from either enzymatic post-translational modification of host targets or misregulation of specific host cell GTPases by *L. pneumophia* proteins.^23^ Loss of individual translocated proteins rarely has negative consequences on intracellular growth, indicating that the system is built with multifold parallel biochemical strategies to ensure intracellular replication.^24^ The importance of coordinating this complex system is emphasized by the central role in disease of the T4SS, which in its most dramatic form results in Legionnaire’s disease, a potentially lethal pneumonia resulting from growth in alveolar macrophages after inhalation of water supplies bearing *Legionella-laden* ameobae.^25^

The Sde family of *L. pneumophila* proteins is a set of four proteins translocated by the T4SS that are required for optimal growth in amoebae,^26^ with biochemical activities subject to extensive regulation by at least three other effectors.^27–32^ Each Sde protein has three domains, shown by either sequence similarity or biochemical analysis to control ubiquitin dynamics (Fig. 1A,B). The N-terminal deubiquitinase (DUB) domain has been demonstrated to show specificity for hydrolyzing K63-linked poly-ubiquitin chains.^15,33,34^ Carboxyterminal to this region are a nucleotidase/phosphodiesterase (NP) domain followed by a mono-ADP-ribosyltransferase (ART)^35^ that collaborate to catalyze phosphoribosylation of ubiquitin (Ub) and atypical ubiquitination through phosphoribosyl linkages (pR-Ub) (Fig. 1A,B).^15,32^ For three of the family members (SdeA, SdeB and SdeC), the ART domain activates ubiquitin by ADP-ribosylation of ubiquitin Arg42 using the cofactor β-nicotinamide adenine dinucleotide (β-NAD). The resulting ADP-ribosylated ubiquitin (ADPr-Ub) provides a substrate for the Sde-family NP domain, which results either in the crosslinking of monophosphoribosylubiquitin (pR-Ub) to target proteins or hydrolysis to release AMP, forming a pR-Ub (Fig. 1A).^36–39^

**Figure 1.**
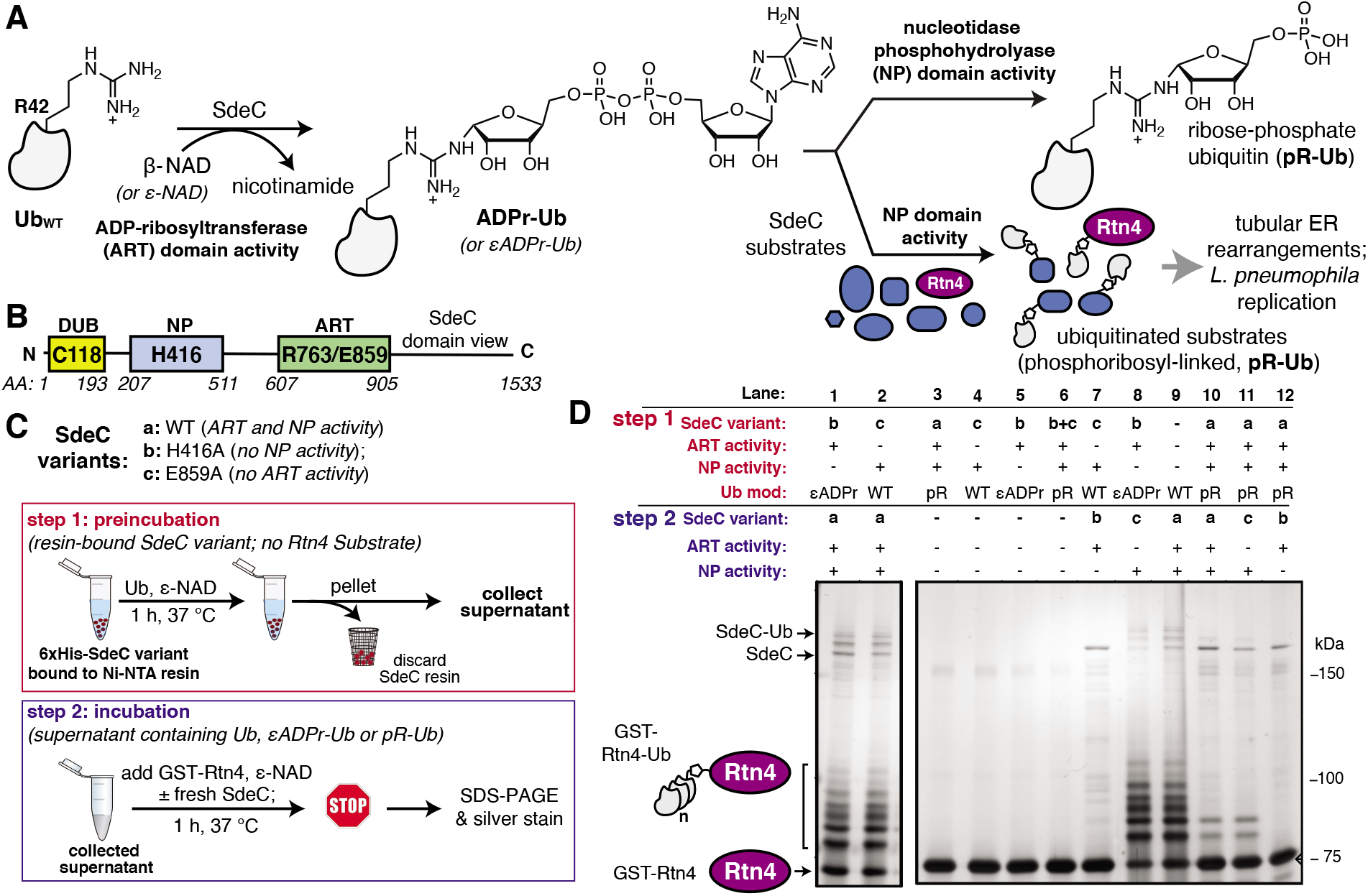
The ART and NP domains of SdeC ubiquitinate substrates in a two-step manner. **A)** Pathway for SdeC-mediated phosphribosyl-linked ubiquitination (pR-Ub).^15^ When monomeric wild-type ubiquitin (Ub_WT_) is incubated with SdeC, the mono-ADP-ribosyltransferase (ART) domain ADP-ribosylates Ub at R42 in the presence of NAD to form ADPr-Ub. Next, the nucleotidase/phosphohydrolyase (NP) domain hydrolyzes AMP from the intermediate and transfers pR-Ub onto the final substrate (bottom path). In the absence of an acceptor substrate, AMP hydrolysis produces phosphoribose-modified ubiquitin (pR-Ub, top path). **B)** Orientation of Sde family protein domains.^35^ **C)** Cartoon scheme depicting the two-step bead assay. In step 1, recombinant purified His-tagged SdeC variants conjugated to Ni-NTA resin were treated with Ub and NAD in the absence of any acceptor substrate. In step 2, the resulting supernatant, which contains unmodified Ub (Ub_WT_), ADPr-Ub or pR-Ub, depending on the SdeC variant used, is incubated with fresh recombinant SdeC and GST-Rtn4 for 1 h at 37 °C. **D)** ADPr-Ub, but not pR-Ub, is a precursor to pR-linked ubiquitination of Rtn4. After treatment using the two-step bead assay, the resulting reaction mixtures are resolved by SDS-PAGE and imaged by silver staining.

A number of host proteins have been identified as pR-Ub acceptors, including endoplasmic reticulum (ER) proteins reticulon 4 (Rtn4)^15^ and LULL1,^28^ the Golgi associated protein GRASP55,^28^ as well as Rab33b and likely other Rab GTPases.^35^ For most of these acceptors, the functional consequence of pR-Ub modification is unclear, although in the case of Rtn4 modification, wholesale ER rearrangements have been demonstrated^15^ while pR-Ub-LULL1 appears to be recruited to the replication vacuole.^28^ Together, these findings support a critical role for pR-Ub in promoting *L. pneumophila* infection in amoebal species.

There has been considerable work exploring the determinants of acceptor protein recognition by Sde family proteins, as pR-Ub modification proceeds *via* a previously uncharacterized pathway distinct from canonical eukaryotic ubiquitination that results in an isopeptide bond linkage between Ub and an acceptor protein lysine.^23,40^ Analysis of model target peptides and crystal structures of the Sde family ART-NP regions have argued that target recognition by the Sde family involves presentation of a Ser residue in unstructured regions of acceptor proteins,^32,37–39^ enabling phosphoribosyl-linked ubiquitination of these cellular targets. This has led to the concept that Sde proteins are Ser-specific pR-Ub transferases. In this manuscript, we demonstrate for the first time that Tyr residues are efficient acceptors of pR-Ub. Using a combination of intact protein and model peptides, we have shown that there is tolerance for a variety of Tyr acceptors with surprisingly little evidence for Ser preference. This result explains previous work consistent with cryptic poly-pR-Ub modification of targets,^15,31^ adding a previously unappreciated level of complexity to the analysis of Sde family activity.

## Results

### Phosphoribosylubiquitination is promoted by SdeC in two independent steps

The crystal structures of the *Legionella* SdeA demonstrate that the ART and NP active sites face in opposite directions and are located 50 Å apart.^37–39,41^ Therefore, it is likely that the two activities work in a nonconcerted fashion. Moreover, the ART and NP domains are known to be biochemically independent, as a functional mutation in one domain has no impact on the catalytic properties of the other. This argues that ADP-ribosylated ubiquitin (ADPr-Ub) does not need to be directly presented to the NP active site by the ART domain, making it likely that the two domains operate as if they were two separable proteins (Fig. 1B). Furthermore, size fractionated ADPr-Ub incubated with SdeA derivatives having catalytically inactive ART domains are able to pR-Ub modify substrates, further emphasizing the independence of these activities.^39,41^ To develop an assay that allows rapid analysis of modified forms of Ub and facile interrogation of the independent activities, agarose bead-immobilized purified recombinant SdeC variants^15^ were used to allow easy separation of enzyme from substrates. This strategy centered around sequential incubation of Ub with a resin-bound SdeC variant followed by removal of the immobilized SdeC (*step 1*), allowing additional immediate treatment of the modified Ub with a different SdeC derivative in the presence of an acceptor protein (*step 2*) (Fig. 1C).

N-His-SdeC variants (see Materials and Methods) were immobilized on nickel agarose resin and the bead-bound enzymes were incubated with recombinant human monomeric Ub and ε-NAD during a “preincubation” step. The immobilized SdeC variants were then removed and the resulting modified ubiquitin species, present in the supernatant, were subsequently treated in an “incubation” step with SdeC variants in the presence of GST-Rtn4, a known Sde-mediated pR-Ub acceptor.^15^ This assay allowed preparation of either ADPr-Ub or pR-Ub, depending on the SdeC variant used,^15^ followed by evaluating which of these isolated species could be crosslinked to GST-Rtn4.

After preincubation in the absence of SdeC (Fig. 1D, Lane 9) or with catalytically dead ART or NP variants of SdeC (Fig. 1D, Lanes 1, 2), subsequent incubation of the resulting modified Ub species with GST-Rtn4, SdeC_WT_, and NAD resulted in the previously reported multiple GST-Rtn4-pR-Ub and SdeC-pR-Ub due to multi-monoubiquitination (Fig. 1D, lane 9).^15,37^ This indicates that the preincubation step does not interfere with subsequent pR-Ub crosslinking. Furthermore, regardless of the SdeC variant used in the preincubation step, GST-Rtn4 cannot be linked to pR-Ub without subsequent incubation with additional SdeC (Fig. 1D, Lanes 3-6). Notably, preincubation of Ub with intact SdeC_WT_ in the absence of acceptor dramatically reduced the amount of Ub available for crosslinking to GST-Rtn4 during the subsequent incubation step (Fig. 1D, Lanes 10-12). As incubation of Ub with SdeC results in pR-Ub,^15^ this indicates that pR-Ub is an end-product that cannot be used for crosslinking to acceptor molecules. Therefore, unlike the traditional Ub pathway, Sde-mediated phosphoribosyl-linked ubiquitination can generate end-products that do not result in productive linkages, yet consume free Ub.

In contrast to the use of pR-Ub as a substrate, the presentation of ADPr-Ub to SdeCE859A, a SdeC variant lacking a functional ART domain, allowed robust pR-Ub crosslinking to GST-Rtn4 that was indistinguishable from incubation with SdeC_WT_ (Fig. 1D, Lanes 8,9). If the order of treatment were reversed, in which Ub was first preincubated with the immobilized SdeCE859A (ART-defective) and then incubated with SdeCH416A (NP-defective), crosslinking to GST-Rtn4 was dramatically reduced (Fig. 1D, Lane 7). Therefore, an ordered series of steps catalyzed by independently acting enzymatic activities is necessary and sufficient for pR-Ub crosslinking to substrate. On the other hand, if SdeC modifies Ub in the absence of substrate, pR-Ub is formed,^15^ acting as a terminal end-product that prevents transfer of Ub to the acceptor protein.

### SdeC catalyzes atypical ubiquitination through tyrosine

In the course of performing preincubations in the presence of hemagglutinin epitope (HA)-tagged Ub, we found that incubation of the tagged derivative with bead-immobilized SdeC_WT_ and NAD resulted in a regular pattern of band laddering, even in the absence of a known substrate (Fig. 2A,B). Although not previously analyzed, these laddering species could be seen in previous work.^15,31^ We performed additional experiments using purified proteins to confirm that this phenomenon is exclusive to HA-Ub and did not occur with monomeric Ub_WT_ or His-tagged Ub (Fig. S1A,B). When treated with either SdeC_E859A_ (ART-defective) or SdeC_H416A_ (NP-defective), no laddering was observed, indicating that crosslinking required functional ART and NP domains (Fig. S1A). When we performed the reactions with ε-NAD, which is a homolog of β-NAD that allows for facile detection of ADP-ribosylation (Fig. S1B), the SdeC_H416A_ (NP-defective)-treated HA-Ub exhibited high levels of ADP-ribosylation, as expected for SdeC variants lacking NP activity. Accordingly, Ub substrates treated with SdeC_WT_ exhibited low levels of ε-ADP-ribosylation, consistent with a crosslinking attack that generates a pR-linkage between Ub and target, with associated liberation of ε-AMP. Therefore, the HA tag provided an unexpected Sde substrate that could be targeted for pR-Ub crosslinking.

**Figure 2.**
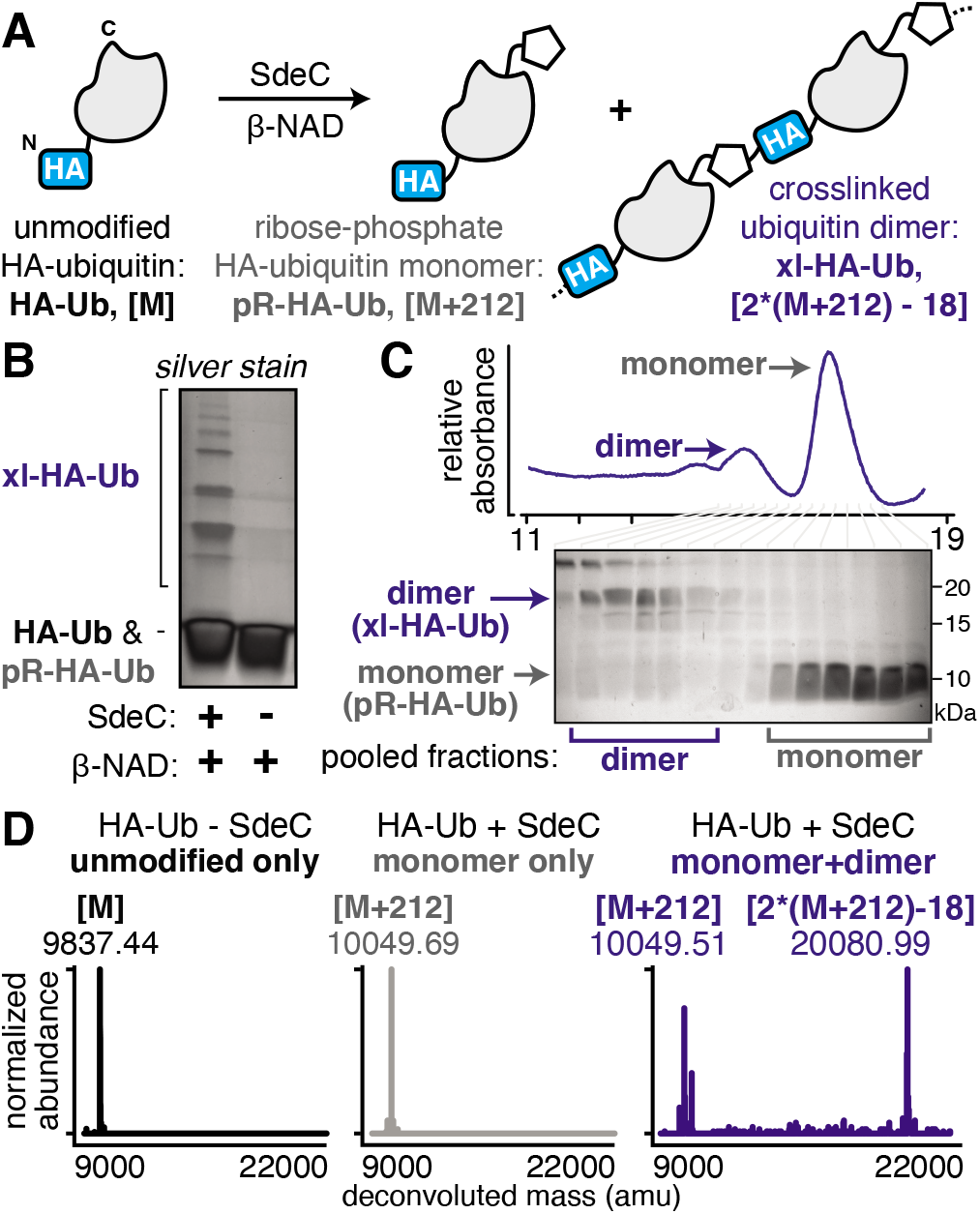
SdeC catalyzes crosslinking of HA-tagged ubiquitin. **A)** Scheme depicting the SdeC-catalyzed phosphoribosylation and/or crosslinking of HA-tagged ubiquitin (HA-Ub) in the presence of β-NAD. **B)** Crosslinked HA-Ub products resolved by SDS-PAGE and visualized by silver stain. **C)** After treatment with SdeC, crosslinked HA-Ub dimers (xl-HA-Ub) were separated from monomeric phosphoribosylated HA-Ub (pR-HA-Ub) using size exclusion chromatography (see Materials and Methods for details). **D)** Pooled fractions of monomeric pR-HA-Ub (gray) or crosslinked products (xl-HA-Ub, blue), as well as untreated HA-Ub (black) were analyzed by deconvoluted protein mass spectrometry. As expected, the monomeric pR-HA-Ub exhibits an [M+212] mass shift relative to the unmodified HA-Ub, consistent with phosphoribosylation. In the pooled dimer fractions, a [2*(M+212)-18] mass shift was observed, which matches with the mass change expected for crosslinking of two pR-HA-Ub monomers with the loss of water. Representative deconvoluted mass spectra for HA-Ub without SdeC treatment (black, [M]), monomeric SdeC-treated HA-Ub (pR-Ub, gray, [M+212), and crosslinked (predominantly dimeric) SdeC-treated HA-Ub (xl-Ub, blue, [(M+212)-18]).

The HA epitope (MYPYDVPDYA) lacks Ser residues that are the supposed targets for phosphoribosyl-linked ubiquitination,^37^ but is the only feature that distinguishes substrates that polymerize (HA-Ub) from those that do not (such as Ub_WT_). Thus, we decided to further investigate this surprising finding to determine if the SdeC-mediated pR-Ub linkage extends beyond modification at Ser only. The Ub sequence in both HA-Ub and Ub_WT_ contains three serine residues, indicating that in the proper context, residues other than Ser are likely targets. We first confirmed that polymerization of HA-Ub occurred through a mechanism analogous to Ub_WT_ crosslinking with Rtn4 (Fig. 2A). To do so, we analyzed HA-Ub treated with or without bead-immobilized SdeC and β-NAD using intact protein liquid chromatography-mass spectrometry (LC-MS). Only a fraction (roughly 10-20%) of the HA-Ub substrate polymerized, so size exclusion chromatography was used to separate monomeric HA-Ub fractions from those that were polymeric after treatment with SdeC and β-NAD (Fig. 2B,C), and the resulting isolated preparation was used for the subsequent analysis.

Based on LC-MS, the pooled monomeric fractions were quantitatively converted to an [M+212] adduct (*observed* 10,049.69 amu, *expected* 10,0049.21 amu) with virtually no unmodified HA-Ub remaining (*observed* 9,837.44 amu, *expected* 9837.21 amu). This is the predicted result for SdeC-mediated formation of phosphoribose-modified HA-Ub (pR-HA-Ub, Fig. 2A,D).^15^ In the pooled polymeric fractions, which were enriched with dimeric species, LC-MS analysis revealed an additional mass of 20,080.99 amu (*expected* 20,0080.42 amu), consistent with crosslinking of two pR-HA-Ub monomers with the loss of water ([2*(M+212)-18]; xl-HA-Ub). Based on the results obtained in Fig. 1 demonstrating a two-step reaction, this was the mass change predicted by a crosslinking mechanism in which ADP-ribosylated HA-Ub provides a substrate for condensation with another pR-HA-Ub monomer, thereby generating xl-HA-Ub (Fig. 2A).^15^

Past work has established that R42 is the site of ADP-ribosylation on ubiquitin.^15,32^ To determine the complementary site of crosslinking in HA-Ub polymers, we subjected the purified monomer (pR-HA-Ub) or dimer (xl-HA-Ub) fractions to proteolysis by trypsin and subsequent LC-MS/MS analysis (Fig. 3A). We began by searching for the phosphoribosylated tryptic fragment (E34-K49 [M+212]: 1882.93 amu) that exists primarily as the doubly and triply charged ions (m/z 940.96 [z=2] & 627.64 [z=3]). These ions were abundant in the monomeric (pR-HA-Ub) sample, but were substantially reduced in the dimeric (xl-HA-Ub) sample (Fig. 3B). This observation is consistent with our previous findings that the intermediate ADPr-HA-Ub is consumed during crosslinking, thereby diminishing the levels of pR-HA-Ub that could be obtained. Based on the observation that crosslinking does not occur in the absence of an HA-tag, we investigated the likelihood that crosslinking occurs between the Ub tryptic fragments that contains R42 (E34-K49) and the fragment that contains the HA tag (M1-K17). The predicted mass of the resulting crosslinked xl-Ub tryptic fragment between R42 of Ub and a site within the HA tag is 3897.84 amu. We were able to confirm the presence of this xl-HA-Ub tryptic fragment exclusively in the dimeric, but not monomeric, samples using an extracted ion chromatogram (EIC) of the triply and quadruply charged ion (m/z 1300.96[z=3] & 975.79[z=4]) (Fig. 3B,C).

**Figure 3.**
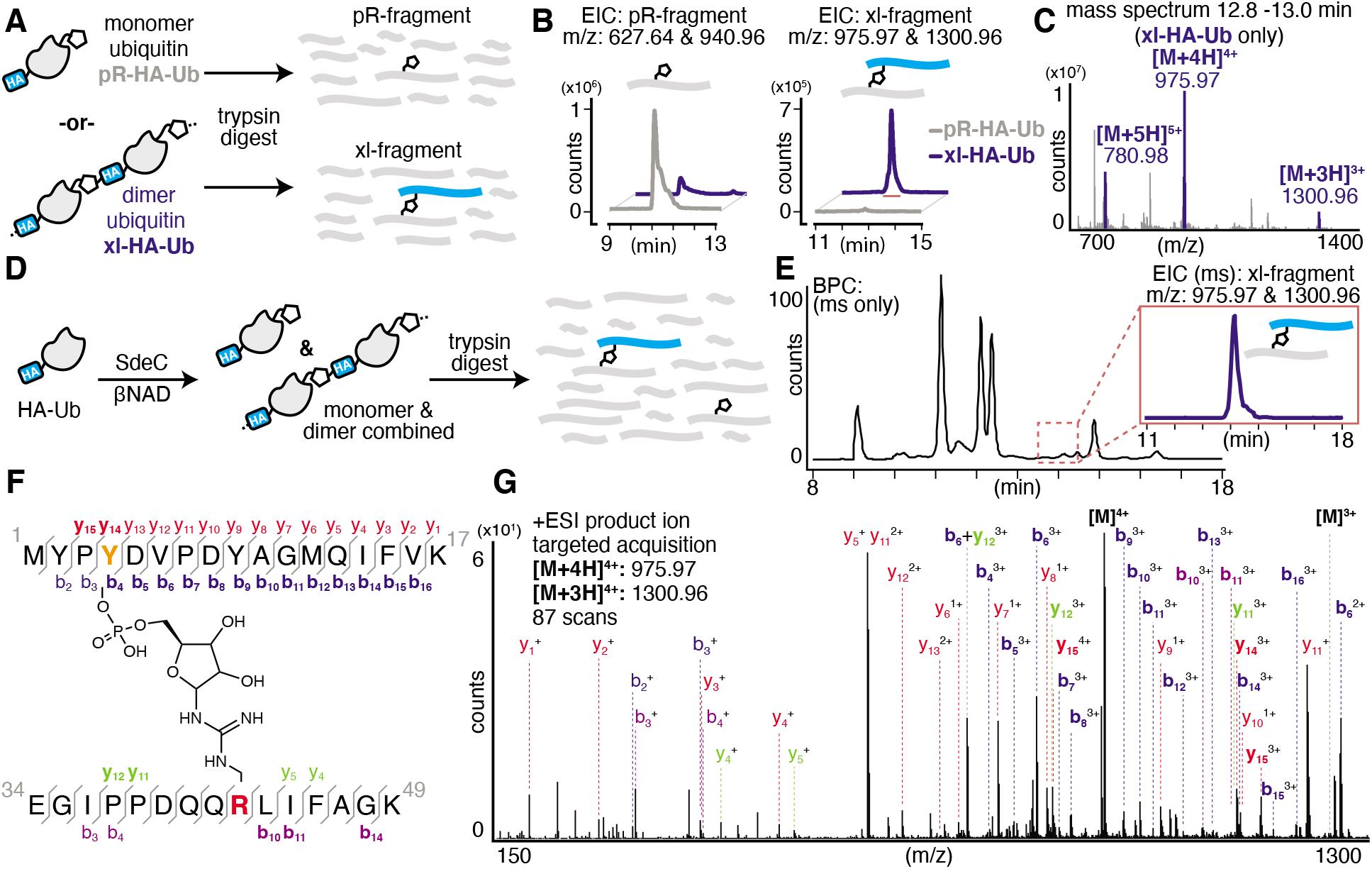
SdeC catalyzes atypical ubiquitination through tyrosine. **A)** Pooled fractions of monomeric phosphoribosylated HA-Ub (pR-HA-Ub, black) or crosslinked products (xl-HA-Ub, blue) subjected to proteolytic digest by trypsin and subsequent analysis by LC-MS. **B)** Extracted ion chromatograms (EIC) for the mass corresponding to the tryptic Ub fragment (pR-fragment, residues 34-49) phosphoribosylated at R42 of HA-Ub and the predicted tryptic Ub fragment (xl-fragment) crosslinked to a residue within the N-terminal HA tag. The mass corresponding to the crosslinked fragment was only observed upon SdeC treatment, and further was only observed for purified dimeric/polymeric (blue) products and was not found in monomeric samples (black). **C)** Combined mass spectrum for the peak corresponding to the tryptic crosslinked fragment in panel **B**. **D)** Scheme depicting the SdeC-catalyzed phosphoribosylation and crosslinking of HA-tagged ubiquitin (HA-Ub) in the presence of β-NAD. The resulting mixture of phosphoribosylated (pR-HA-Ub) and crosslinked (xl-HA-Ub) HA-Ub was subjected to proteolytic digest by trypsin and subsequent analysis by LC-MS/MS. **E)** Representative base peak chromatogram for the SdeC-catalyzed phosphoribosylation and crosslinking of HA-tagged ubiquitin. The crosslinked tryptic fragment (residues 1-17) could be readily identified using an extracted ion chromatogram (EIC) for m/z 975.97 [z=4] & 1300.96 [z=3], which matches with the predicted mass for the xl-fragment. **F)** Subsequent MS/MS analysis showing Y4 crosslinking, (out of three possible tyrosine residues in the HA sequence). **G)** Relevant b and y ions identified for each of the tryptic fragments are indicated. Observed ions that are specific only to modified (but not unmodified) peptides are shown in bold *blue*, b ions; *red*, y ions. Ions labeled in purple and green represent MS/MS fragmentation for b or y ions, respectively, derived from the ubiquitin side (rather than HA side) of the crosslink.

Having confidently determined the correct crosslinked tryptic fragment we sought to identify the site of crosslinking within the HA tag. We treated HA-Ub with SdeC and β-NAD and subjected the entire sample to proteolysis by trypsin without any further purification by size exclusion chromatography to avoid sample dilution (Fig. 3D). Using this approach, we could readily identify the xl-HA-Ub tryptic fragment by extracted ion chromatogram (Fig. 3E). MS/MS analysis of this xl-HA-Ub trypic fragment enabled us to determine that crosslinking occurs at Y4 within the HA sequence, as indicated by the identified b and y ions (Fig. 3F). Observation of modified b4 and y14 ions, which describe xl-HA-Ub ions fragmented at the amide just after or just before Y4, respectively, allowed us to unambiguously conclude that the site of crosslinking for the HA-Ub protein is at Y4, despite the presence of two additional tyrosine residues within the HA sequence (Fig. 3F,G and Table S1). Thus, these results establish that Sde family proteins can catalyze phosphoribosyl-linked ubiquitination at tyrosine, in addition to serine.

### SdeC is promiscuous with respect to crosslink formation

The serendipitous discovery of the HA epitope as an acceptor substrate for phosphoribosyl-linked ubiquitination offered a unique opportunity to explore the specificity of Sde family proteins. Although there are three tyrosines in the HA sequence, we found that only one (Y4) was modified when the HA-tag was fused to a full-length protein like ubiquitin. To further corroborate the observed site selectivity of SdeC, and to determine if there is a molecular basis for this preference, we used a series of synthetic HA peptide variants that substituted Phe in place of Tyr. In doing so, the ability of the HA peptide to be a crosslinking acceptor was evaluated one Tyr at a time (HA_Y2F_, HA_Y4F_, HA_Y9F_), and then all together (HA_Y249F_). We evaluated these variants, as well as the wild-type HA peptide (HA_WT_), as potential SdeC substrates using a combination of SDS-PAGE and intact protein LC-MS. SdeC was still capable of catalyzing phosphoribosyl-linked ubiquitination on the wild-type HA peptide when co-incubated with monomeric Ub and β-NAD (Fig. 4A,B). Surprisingly, each single point variant (HA_Y2F_, HA_Y4F_, HA_Y9F_) led to appreciable levels of crosslinked product that were observable by SDS-PAGE and intact LC-MS, including HA_Y4F_ (Fig. 4B,C), which would be expected to be a poor acceptor based on the preference for Y4 using the HA-Ub substrate (Fig. 3). In only the triple variant, in which all three potential tyrosine acceptors are removed, is there a complete loss of crosslinking (Fig. 4B,C). Therefore, although there is selectivity for the exact site through which polymerization occurs for full-length HA-Ub proteins, there appears to be promiscuity when peptide HA substrates are used.

**Figure 4.**
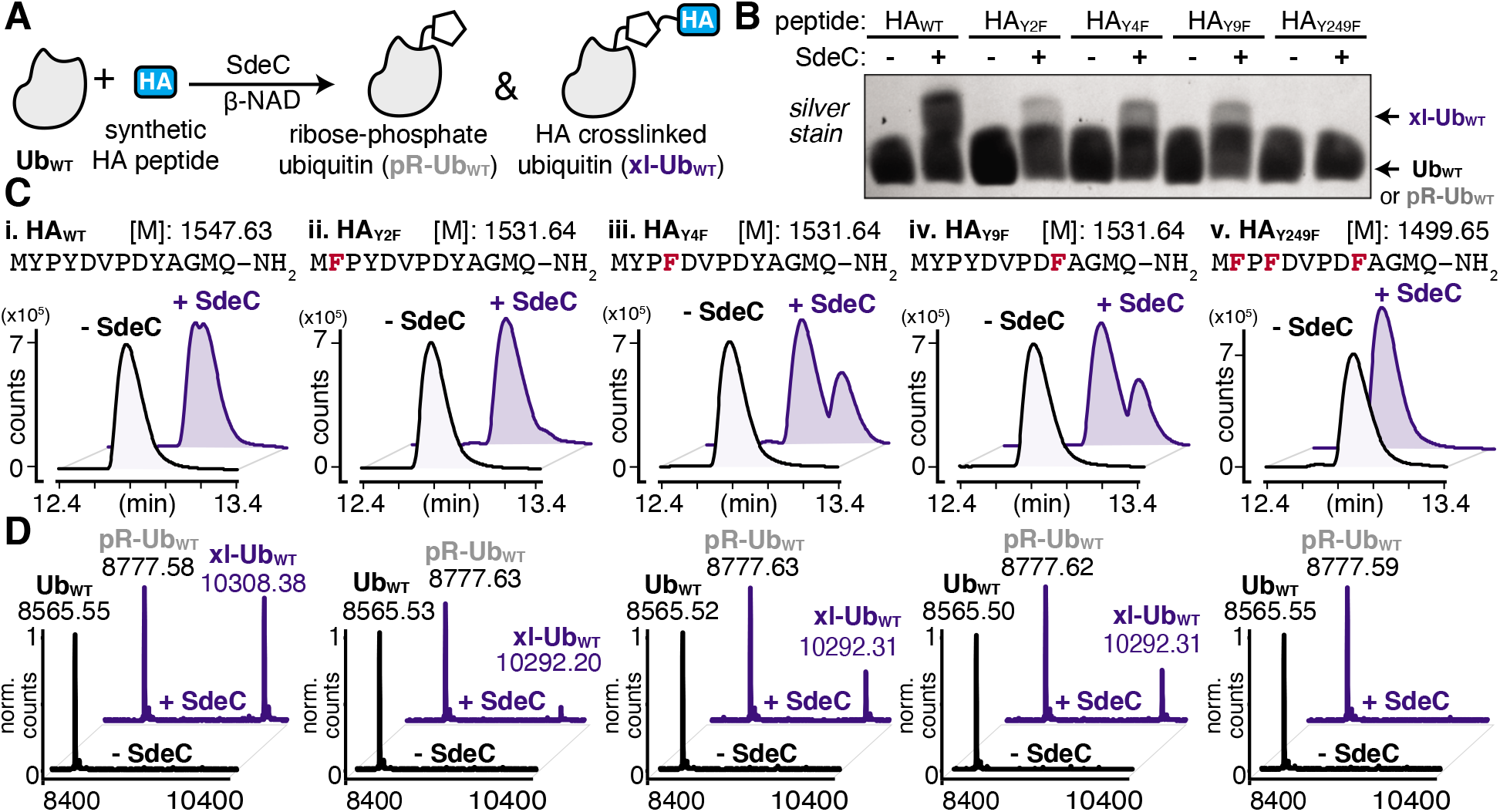
SdeC can utilize synthetic HA peptides as substrates for crosslinking. **A)** Scheme depicting the SdeC-catalyzed phosphoribosylation of wild-type ubiquitin (Ub_WT_) and its crosslinking to synthetic HA peptides in the presence of β-NAD. **B)** Crosslinking of synthetic HA peptide variants in which Tyr is substituted with Phe, assessed by SDS-PAGE and silver staining. **C)** HA_WT_, as well as HA_Y2F_, HA_Y4F_, HA_Y9F_ and HA_Y249F_ peptide variants were incubated as in (A) and analyzed by intact protein mass spectrometry. Representative base peak chromatograms and **(D)** deconvoluted mass spectra for Ub_WT_ alone (black), or treated with SdeC (blue) and the indicated synthetic HA variant.

Based on the surprisingly persistent crosslinking for HA point variants, we next sought to determine if crosslinking could occur at multiple Tyr residues. Proteolysis by trypsin followed by LC-MS/MS analysis identified tryptic fragments corresponding to HA peptides crosslinked with full-length Ub (Fig. 5A,B). Notably, the abundance of the observed crosslinked fragment for each of mutant derivatives was reduced compared to the HA_WT_ peptide (Fig. 5B). These results are consistent with a model in which each tyrosine residue can act as an acceptor. Therefore, SdeC is able to use any available Tyr residue located on short unstructured peptide substrates as the final acceptors.

**Figure 5.**
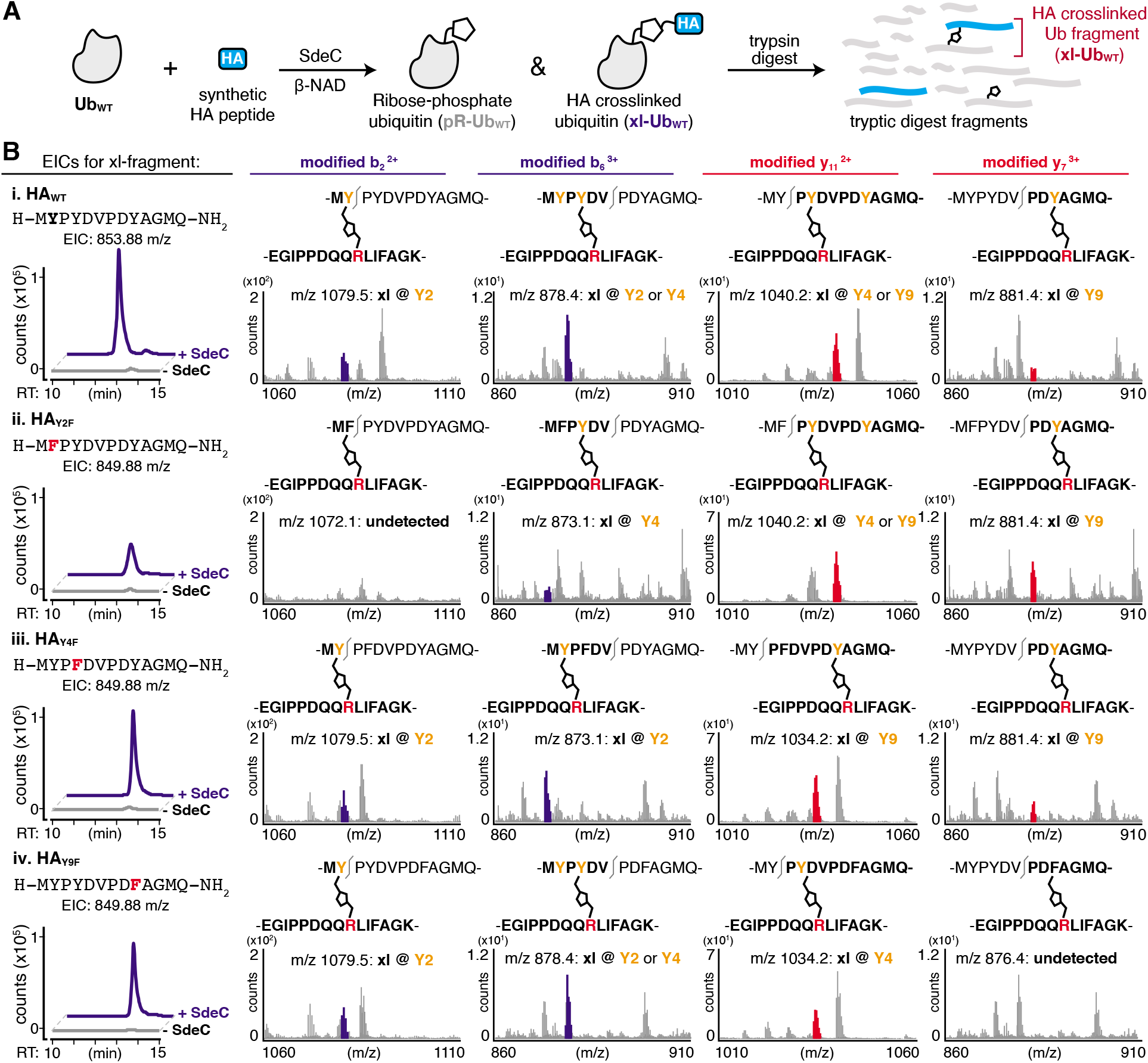
SdeC is promiscuous with respect to crosslink formation at Tyr. **A)** Scheme depicting the SdeC-catalyzed phosphoribosylation of ubiquitin (Ub_WT_) and its crosslinking to synthetic HA peptides in the presence of β-NAD. **B)** After tryptic digest, peptide mixtures were analyzed using LC-MS/MS. Displayed are extracted ion chromatograms (EICs) for the mass-to-charge ratio (m/z) of fragments corresponding to synthetic HA peptide crosslinked to Ub_WT_ (xl-Ub_WT_). **C)** Crosslinking occurs at each of three Tyr residues in synthetic peptides. Key diagnostic fragmentation ions are highlighted. Collectively, these ions indicate that synthetic HA peptide is able to crosslink *via* each of the three tyrosine residues in its sequence.

We next determined if more than one site in a single peptide could act as an acceptor. In pursuing this hypothesis, we searched for diagnostic b or y ions that could confirm that the xl-Ub tryptic fragment (Fig. 5B, Table S2) was not a single species but rather a mixture of xl-Ub isomers, each of which formed a crosslink at a distinct single site. For instance, the identification of a modified b2 ion confirms crosslinking at Y2. Such a b_2_ ion was identified for crosslinked HA_WT_, HA_Y4F_ and HA_Y9F_ peptides, but was absent in HA_Y2F_ (Fig. 5C, left). Similarly, the identification of a modified y7 ion is diagnostic of crosslinking at Y9, and was present in tryptic fragments generated from HA_WT_, HA_Y2F_ and HA_Y4F_ peptide variants, but was absent from HA_Y9F_ (Fig. 5C, right). Crosslinking at Y4 generated a modified y_11_ ion in the HA_Y9F_ variant while it resulted in a modified b6 ion in the HA_Y2F_ variant (Fig. 5C, middle). These results further support the observation that SdeC catalyzes phosphoribosyl-linked ubiquitination at Tyr in addition to Ser. Therefore, the chemical identity of the acceptor side chain is not limited to Ser as previously suggested, and target specificity by SdeC is promiscuous with respect to the exact site of modification.

### SdeC does not have an intrinsic preference for Ser over Tyr linkages

Having established that SdeC is capable of catalyzing crosslinking to Tyr in addition to Ser, we next sought to evaluate if there were a preference for a particular hydroxyl sidechain. To evaluate the relative residue preference of phosphoribosyl-linked ubiquitination, we prepared a synthetic peptide variant that substituted all tyrosines for serines (HA_Y249S)_. After treatment of monomeric Ub and HA_Y249S_ with SdeC and β-NAD, the levels of crosslinking to HA_Y249S_ were found to be comparable to incubation with HA_WT_, consistent with previous reports that Ser can act as a substrate for SdeC (Fig. 6A,B). To analyze if there were preferences for the chemical nature of the sidechain, a panel of single amino acid variants that individually replaced each Tyr with Ser were evaluated for crosslinking (HA_Y2S_, HA_Y4S_ & HA_Y9S_; Fig. 6B). Individual replacement of each Ser residue maintained crosslinking to the three peptides (Fig. 6B,C). Based on these results, the nature of residue preference was analyzed further.

**Figure 6.**
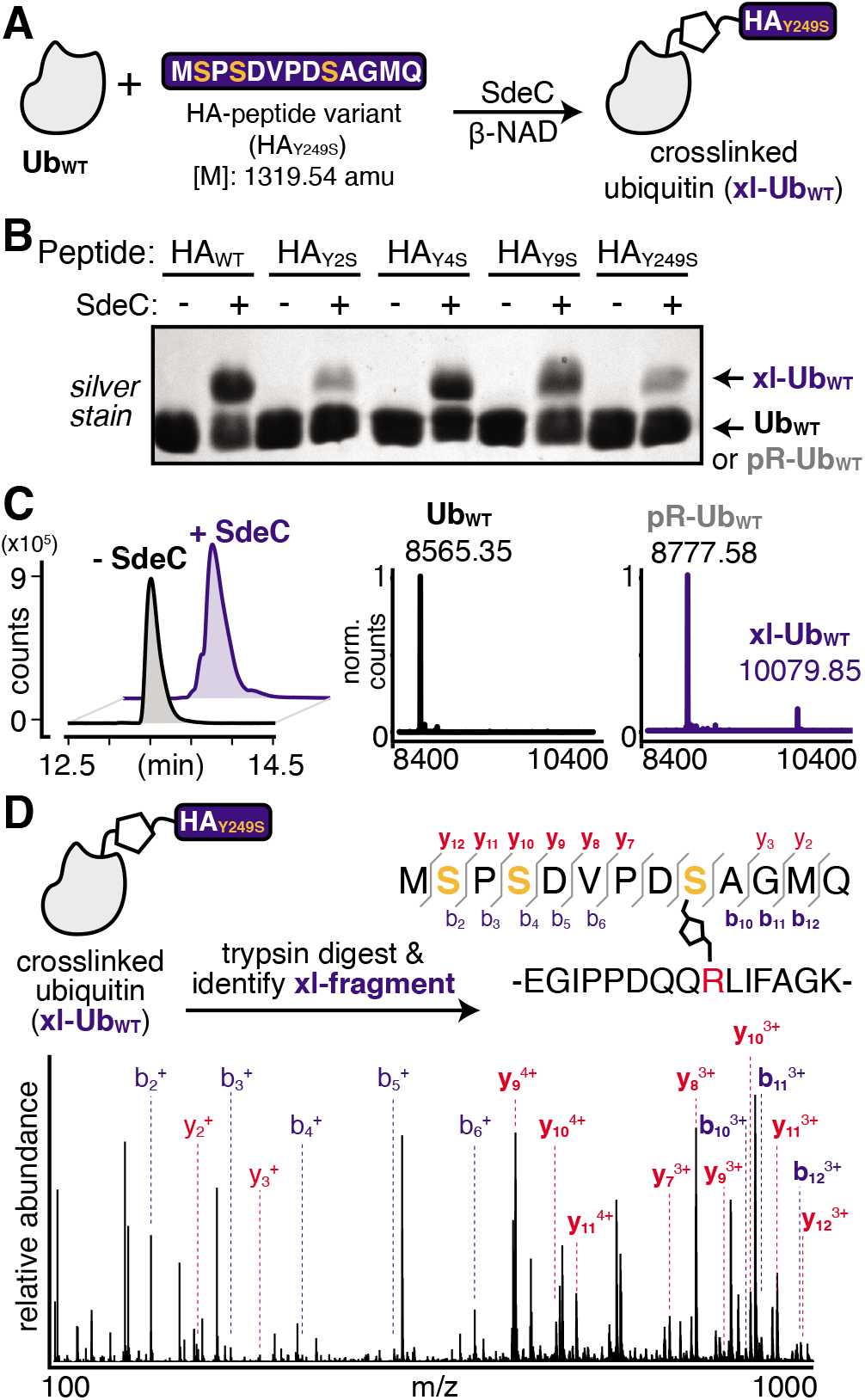
The observed site selectivity for phosphoribosyl-linked ubiquitination is dependent on the hydroxyl substrate. A) Scheme depicting the reaction of Ub_WT_ with a triple serine variant (HA_Y249S_). The full sequence is shown for clarity. **B)** SdeC is able to catalyze the crosslinking of this serine variant to Ub_WT_, as determined by intact protein LC-MS. Base peak chromatograms (BPCs, left) show a shift upon treatment with SdeC. Deconvoluted mass spectra (right) reveal that no reaction has occurred in the absence of SdeC (black), but upon SdeC treatment (blue) Ub_WT_ is quantitatively phosphoribosylated ([M+212] = 8777.58), and partially undergoes crosslinking with HA_Y249S_ (xl-Ub_WT_: 10079.85). **C)** MS/MS analysis confirms that crosslinking occurs at S9 in HA_Y249S_. The protein mixture was digested using trypsin and the crosslinked tryptic fragment was identified (exact mass: 3185.48 amu; m/z: 796.62 [z=4] and 1061.82 [z=3]. Relevant b and y ions identified for each of the tryptic fragments are indicated. Observed ions that are specific only to modified (but not unmodified) peptides are shown in bold *blue*, b ions; *red*, y ions.

MS/MS analysis of crosslinking to HA_Y249S_ found that crosslinking to the triple-Ser variant shifted the position preference to S9 (Fig. 6D, Table S3). MS/MS analysis of single Ser variants supported this result, as any convincing evidence for crosslinking to serine was limited to S9, as observed for the HA_Y9S_ peptide (Fig. S2, Table S4). There was no indication of crosslinking to S2 (HA_Y2S_) and evidence for crosslinking to S4 (HA_Y4S_) was inconclusive as it could also be attributed to modification at Y2 or Y9. Surprisingly, in all single serine variants, crosslinking to Tyr appeared to remain efficient (Fig S2). For instance, SdeC treatment of the HA_Y9S_ mutant led to robust crosslinking at Y2 and Y4 (Fig. S2, bottom panels). These data argue against an absolute preference for Ser as a target of pR-Ub modification, and are consistent with the surrounding protein microenvironment determining SdeC target site preference.

## Discussion

Previous work has shown that the *Legionella* T4SS effector Sde family catalyzes a non-canonical two-step phosphoribosyl-linked ubiquitination reaction that is completely independent of the host ubiquitination machinery.^15,32,35^ In the absence of an acceptor protein, water can act as a hydroxyl donor, resulting in phosphoribosyl-modifed Ub (pR-Ub) at the Arg42 residue. Using a rapid assay in which immobilized SdeC derivatives can be sequentially incubated with various combinations of substrates, we showed that pR-Ub is a terminal product that cannot be transferred to target proteins. Although, by consensus, the phosphoribosyl-linked ubiquitination of acceptor molecules has been termed serine ubiquitination,^36,37,42^ here we present evidence that this is incorrect, and that tyrosine can also act as a pR-Ub acceptor in the crosslinking step promoted by the SdeC NP domain (Fig. 1). Previous studies using model peptides as acceptors for pR-Ub modification failed to include Tyr as a potential replacement for Ser, providing an explanation for why this target was overlooked.^37^ These results emphasize the fact that although multiple structural studies of SdeA have been reported, the determinants of motif recognition for pR-linked ubiquitination are still rather obscure.^37–39,41^ The model that unstructured regions of proteins are targets for this modification event is clearly supported by our data, as we observed a switch in substrate preference moving from structured to unstructured targets. Our results further argue, however, that the chemical identity of the acceptor residue also leads to a previously unappreciated level of promiscuity.

The crystal structures of SdeA have revealed that the active sites of ART and NP domains are 90° opposed, making it unlikely that ADP-ribosylation of Ub and transfer to an acceptor molecule are coincident events. Our results further support the model that the two domains are biochemically independent and can be fully uncoupled. The results from the sequential bead-based assay, demonstrating that ADPr-Ub provides a substrate for linkage to Rtn4 by the SdeC NP domain, suggest a strategy that can maximize the number of potential substrates for pR-Ub crosslinking. By having duties assigned to different domains acting independently, this raises the possibility that two different *L. pneumophila* proteins can collaborate in the process during intracellular growth. Biochemically, the SdeC ART domain is known to cause rapid formation of ADPr-Ub, with near complete modification occurring even during short incubations on ice.^15^ Physiologically, this could drive the accumulation of a large pool of ADPr-Ub about the *Legionella* replication vacuole. This pool could be used by the NP domain on the same protein, but there are also several *Legionella* translocated proteins consisting of solitary orphan NP domains that could transfer pR-Ub to an alternative spectrum of substrates.^39,43^ As a consequence, catalytic domain collaboration potentially takes place both within or between proteins, with the latter strategy increasing the diversity of pR-Ub crosslinking targets.

The discovery of tyrosine as a Sde protein modification sites is significant for multiple reasons. From a purely technical aspect, this represents an overlooked modification that has likely gone undetected using biochemical strategies that focus on serine modifications. In regards to the intracellular lifestyle, pR-Ub crosslinking is likely part of an “act quickly, leave abruptly” strategy for *L. pneumophila*. Members of the Sde family are secreted immediately on contact with host cells, ensuring that execution of their activities occurs rapidly.^44^ Shortly thereafter, Sde proteins become inactivated by SidJ ^42,44^ and pR-Ub crosslinking on target proteins is reversed by pR-linkage specific deubiquitinases.^27,28^ Therefore, there is only a short window of time for Sde action. Broadening the target selection to multiple hydroxylated sidechains, including Tyr in addition to Ser, should act in concert with having independently acting domains, as both properties broaden the spectrum of proteins that can act as acceptors in a very short timespan.

This work uncovered new levels of complexity regarding the determinants of pR-Ub target recognition. The unstructured nature of the HA peptide used in these studies supports previous hypotheses that an unstructured targeting site likely provides an optimal conformational fit to the enzymatic surface of the NP domain.^37–39,41^ In the case of targeting of the Tyr hydroxyl, however, we found that the context of the peptide was important in regards to residue site-specificity. When targeting the HA-Ub hybrid, there was specificity for pR-Ub linkage to the Y4 residue (Fig. 3). When a free synthetic peptide was analyzed, this specificity was lost, as peptides having single Tyr to Phe substitutions still retained pR-Ub linkage (Fig. 4). In support of promiscuous site selectivity in unstructured peptide targets, we found clear evidence for linkage to all three Tyr residues based on MS/MS analyses (Fig. 5). Although the HA sequence is likely to be unstructured in all contexts, being linked to a larger protein structure such as Ub results in conformational restraints not observed for free peptides in solution, changing the nature of the recognition event. A free peptide may be threaded into the NP enzymatic site in more than one direction, allowing free access to all the Tyr residues. In contrast, HA-Ub may provide restraints that limit the residues accessible to the site of pR-Ub transfer, conferring residue site selectivity.

The observed selectivity was highly dependent on the chemical nature of the sidechain. Ser-containing HA peptides behaved differently from those with Tyr, as they retained specificity for crosslinking at a single position (S9) (Fig. 6). For the targeting of Ser residues, there are likely to be sequence determinants extending beyond conformational restraints that drive residue site specificity. Modification of Tyr residues, in contrast, showed a high level of promiscuity. This raises the possibility that a protein region compatible with recognition by the Sde NP domain supports increased site promiscuity for Tyr targeting relative to that observed for Ser. Targeting a particular substrate residue with high specificity could be a critical element in determining if a modification generates a new function, as opposed to causing protein dysfunction.

In summary, our work provides powerful evidence that the targeting of the Sde family is much broader than previously hypothesized, with the potential that modification could result in either acquiring new functions or blocking activities depending on the site of modification and its relative promiscuity. It also provides insights on the molecular mechanisms of this noncanonical ubiquitination pathway, as well as potential information for understanding how pathogens control host cellular functions.

## Experimental

### Materials & Methods

Please see the Electronic Supplementary Information for complete details about the Materials and Methods used. Abridged experimental protocols follow in this section.

#### Protein purification and peptide synthesis

SdeC variants were purified as described previously^15,45^. Purified GST-Rtn4 was purchased from MRCPPU. Human recombinant monomeric Ub and HA-Ub were purchased from Boston Biochem (R&D systems). All HA derivative peptides were synthesized in Tufts University Core Facility. For further details about protein expression and purification, please see the Electronic Supplementary Information.

#### Rtn4 Ubiquitination using two-step SdeC bead assay

*In vitro* Rtn4 ubiquitination assays were described previously.^15^ For linking recombinant SdeC derivatives to agarose beads, Ni-NTA agarose resin (Thermo Fisher) was washed 3X with ART buffer and mixed with His6-SdeC followed by incubation for 1 h at 4 °C. Then the SdeC-bound agarose resin was washed 3X with 1X ART buffer before processing to the two-step bead assay. During preincubation, SdeC-bound agarose beads were added to 10 mM Ub, and 100 mM ε-NAD in 1X ART buffer (SdeC final concentration 50nM), and incubated for 1 hr at 37°C. After incubation, the beads were removed from reaction by spin filter columns. The supernatant containing modified Ub and excess NAD were mixed with 400nM GST-HA-Rtn4 and fresh unbound SdeC variants, then incubated for another hour at 37°C. Reaction were terminated by addition of reducing loading buffer and boiling.

#### HA-Ub polymerization assay and HA peptide ubiquitination assays

For HA-Ub polymerization assays, 1 μM HA-Ub was incubated with 20 nM recombinant SdeC and 100 μM Nicotinamide 1, N6-ethenoadenine dinucleotide (ε-NAD, Sigma) or β-Nicotinamide adenine dinucleotide (β-NAD, Santa Cruz Biotech/Sigma) at 37 °C in 1X ART buffer for 2 hours.

For HA peptide ubiquitination assays, 12.5 μM HA peptides were mixed with 10 μM monomeric Ub, and incubated with 20 nM SdeC and 250 μM β-NAD at 37°C in 1X ART buffer for 1 hours. All reactions for gel analysis were terminated by either addition of reducing loading buffer and boiling, or flash freezing in liquid N2 for storage prior to further analysis. For samples analyzed by mass spectrometry (MS), excess salts and NAD were removed *via* buffer exchange, using either Illustra NAP-5 Columns (Cytiva Life sciences) or Amicon Ultra Centrifuge filters, 3K MWCO (MilliporeSigma), eluting in water.

#### FPLC purification and Size Exclusion Chomotography

After crosslinking reactions were complete, samples were concentrated 10X by lyophilization (Labconco Freezone12), then purified FPLC with 1X ART buffer on a Superdex 200 10/300 GL column (flow rate 0.25 ml/min). 0.2 ml fractions from the HA-Ub monomer and dimer peaks were collected and combined based on SDS PAGE fractionation and silver staining.

#### General Protocol for Trypsin Digestion

Using Amicon^®^ Ultra centrifugal filters (Millipore Sigma), intact protein reactions were concentrated and underwent buffer exchange into water, and were diluted into 1:1 t-butanol:water. DTT was added to a final concentration of 3 mM. Reactions were then heated to 65 °C for 35 minutes and subsequently cooled to room temperature. Sequencing grade modified trypsin (Promega) was added for a final 1:25 protease:protein ratio. This digest solution was incubated at 37 °C overnight. The resulting solution of peptides was subjected to LC-MS/MS analysis without further purification.

#### Mass Spectrometry

For intact protein analysis, samples were injected onto a ZORBAX 300SB-C8 column (2.1 x 100 mm, Agilent) and eluted with a water:acetonitrile gradient mobile phase with 0.1% formic acid (0.400 mL/min; 5% −80% over 24 min). The mass spectrometer was utilized in positive mode with a dual electrospray ionization (ESI) source. MS spectra were acquired using the following settings: ESI capillary voltage, 4500 V; fragmenter, 250 V; gas temperature, 350°C; gas rate, 12.5 L/min; nebulizer, 50 psig. Data was acquired at a rate = 5 spectra per second and scan range of 100 – 3000 m/z. Analysis was completed using Agilent MassHunter Bioconfirm (v. B.06.00).

Following tryptic digestion, peptide fragments were injected onto an AdvanceBio Peptide 2.7 μm column (2.1 x 150 mm, Agilent) and were eluted with the same gradient and flow rate as indicated above. MS spectra were obtained using the same parameters as described above. MS/MS spectra were acquired using the following settings: ESI capillary voltage, 4000 V; fragmentor, 150 V; gas temperature, 325°C; gas rate 12 L/min; nebulizer, 40 psig. MS/MS was acquired at 1 spectra per second with a mass range of 300 – 3000 m/z. For all MS/MS acquisitions, targeted precursor ion selection, coupled with set collision energies were used. Full length precursor ion m/z’s were calculated using ChemDraw. Precursors were identified *via* MS scans with the following stringency: medium isolation width, z=2-4, RT = 10 min, ΔRT = 2 min). After identification, precursor ions were subjected to iterative rounds of collision induced dissociation in the collision chamber and subsequent mass identification. MS/MS cycles scanned five ionization energies from 20-28 V, with a 2 V step between scans to sample various ionization events for each precursor ion. Following acquisition, all MS/MS spectra were combined for display. On average, targeted MS/MS acquisition yielded 80-140 spectra for each defined precursor ion. Agilent MassHunter Bioconfirm (v. B.07.00) was used for generation of intact protein deconvolutions. Bioconfirm was also used to generate extracted ion chromatograms (EIC) and additional mass spectra. Posttranslational modifications (PTMs) were identified using PEAKS Studio (v. 7.5) software. For further details about MS and MS/MS acquisition and analysis methods, please see the Electronic Supplementary Information.

## Acknowledgements

This work was supported by NIAID grants R01AI11321 to RRI and R01AI146245 to RRI and RAS. We would like to thank Michael Berne for rapid synthesis of synthetic peptides at the Tufts University Core Facility (TUCF).

---

**Figure S1.**
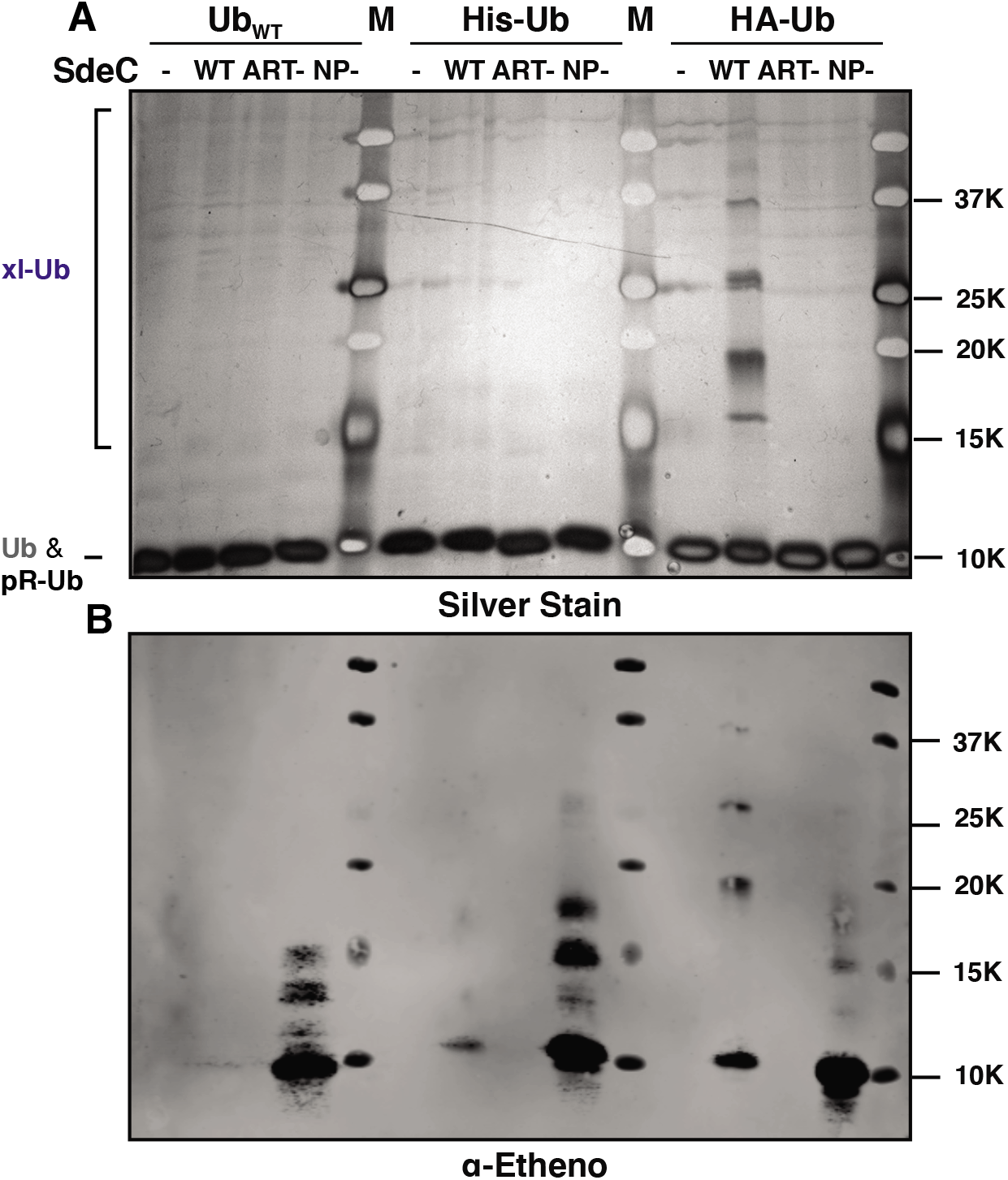
HA-Ub polymerization is dependent on the HA epitope. **A)** To evaluate the polymerization of Ub, Ub_WT_, His-Ub, or HA-Ub were incubated with ε-NAD and SdeC variants at 37 °C for 2 h, then visualized by SDS-PAGE gel and silver staining. **B)** Samples were also evaluated using an α-ethenoadenosine Western blot.

**Table S1.**
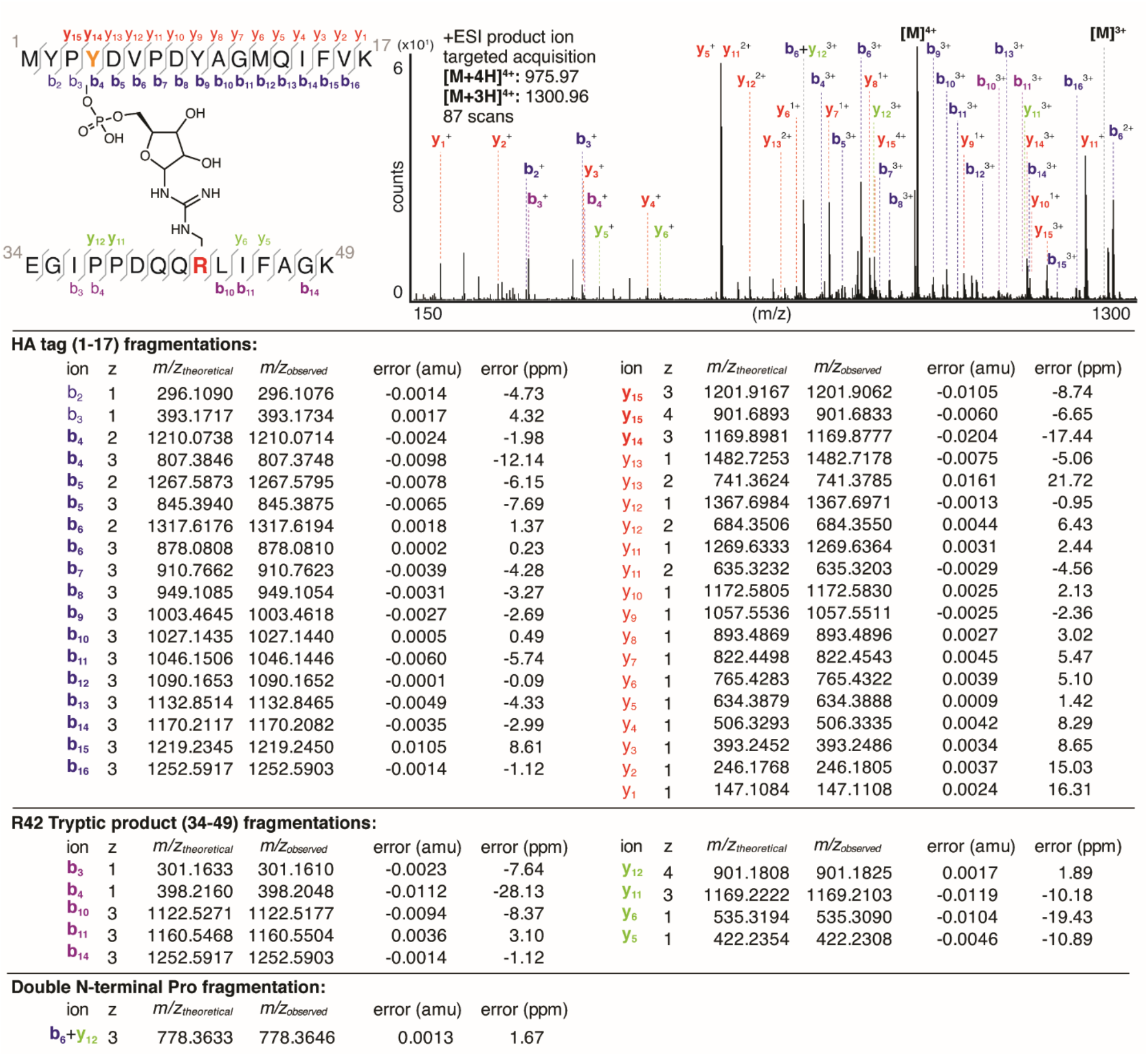
Mass Error Anlysis for HA-Ub xl-fragment.

**Table S2.**
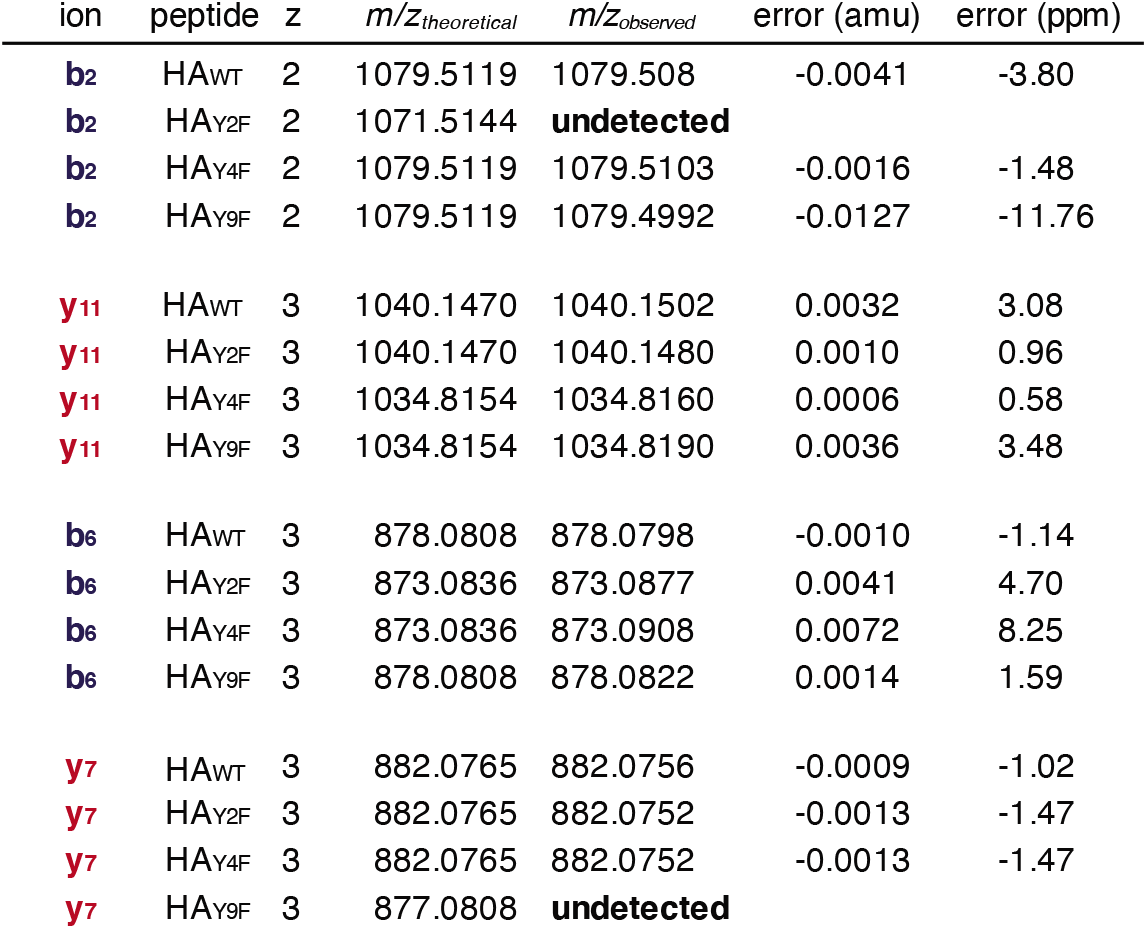
Mass Error Anlysis for Ub_WT_ + HA Peptide Phe-Variants xl-fragment.

**Table S3.**
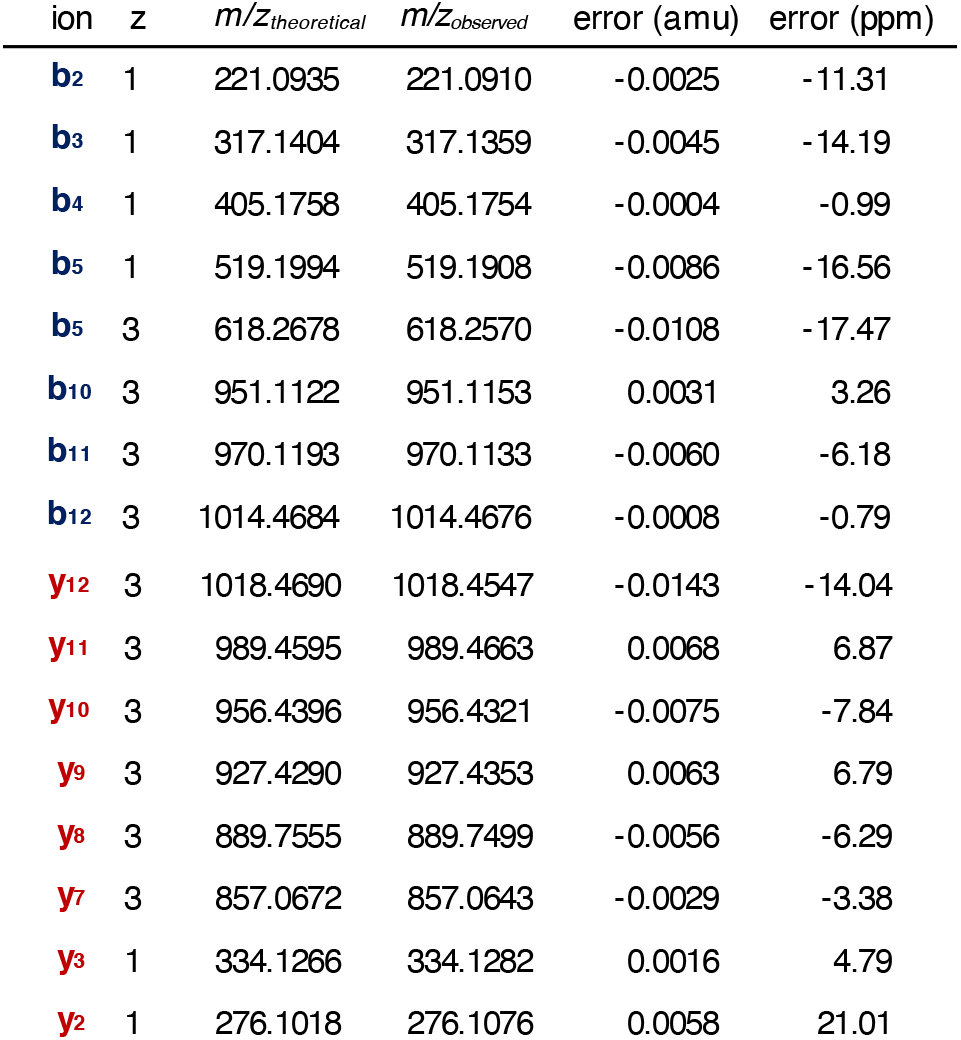
Mass Error Analysis for Ub_WT_ + HAY_249S_ xl-fragment.

**Figure S2.**
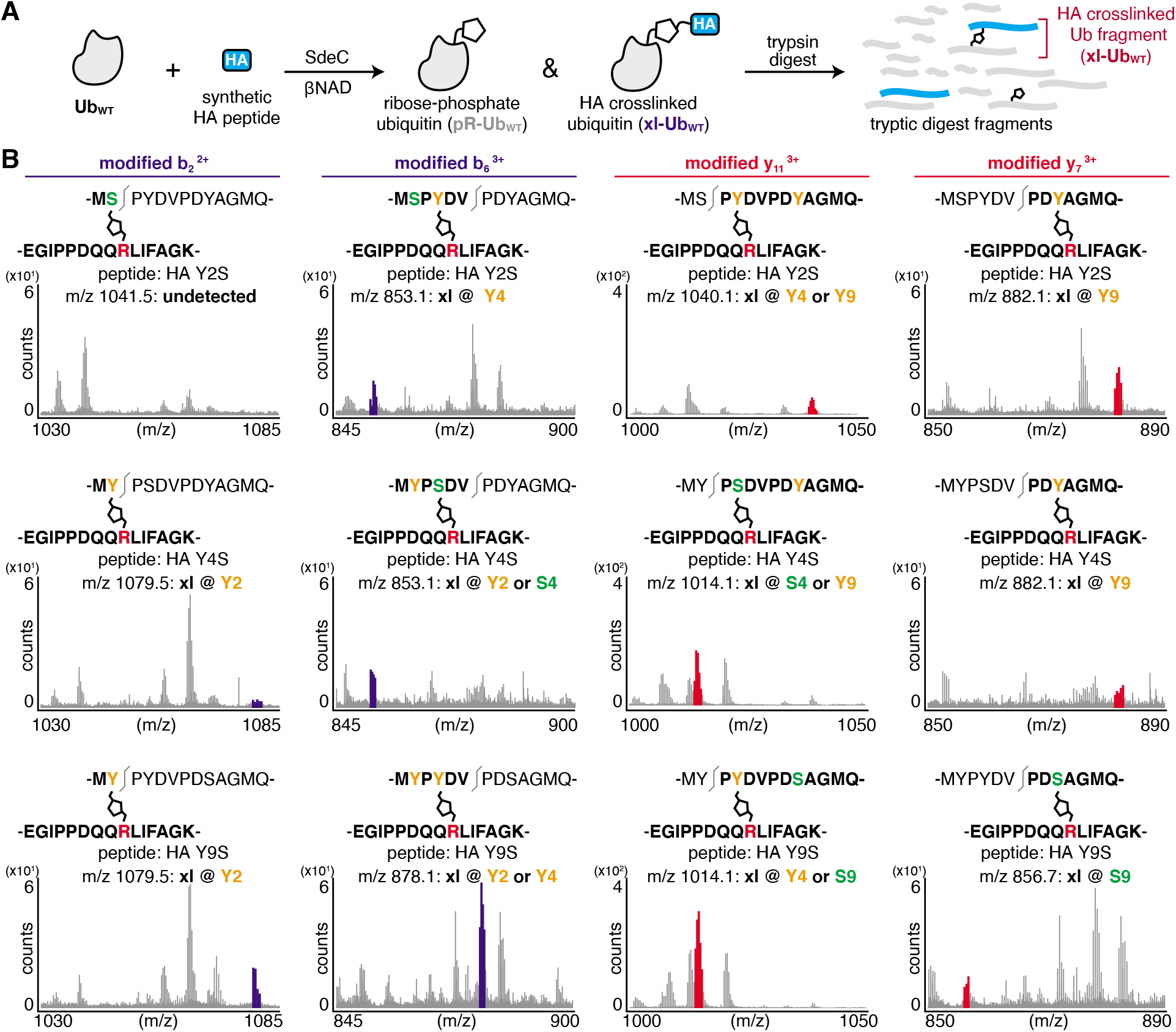
SdeC does not have an intrinsic preference for Ser over Tyr linkages. **A)** Scheme depicting the SdeC-catalyzed phosphoribosylation of ubiquitin (Ub_WT_) and its crosslinking to synthetic HA peptides in the presence of β-NAD. **B)** After tryptic digest, peptide mixtures were analyzed using LC-MS/MS. Key, diagnostic fragmentation ions are highlighted, indicating that crosslinking can occur at multiple positions within the HA peptide. The m/z for the modified b2 ion, which would suggest crosslinking at Y2 or S2, is absent in HA_Y2S_, but present in HA_Y4S_ and HA_Y9S_. The modified b_6_ ion is present in all three peptide variants, and indicates crosslinking either at Y4 for HA_Y2S_. For HA_Y4S_, the presence of this ion suggests that the site of modification may be at either Y2 or S4. For HA_Y4S_, the presence of this ion suggests that the site of modification may be at either Y2 or Y4. Similarly, a modified y_11_ ion is present in all peptide variants, though this ion would suggest two possible sites of crosslinking, each highlighted in orange, as indicated for each spectra. Finally, the modified y_7_ ion indicates crosslinking at Y9 or S9 only. We found that the only definitive evidence of modification at serine for these peptides occurred at S9 for HA_Y9S_. For HA_Y2S_ and HA_Y4S_, this ion is present and indicates that crosslinking occurs at Y9.

**Table S4.**
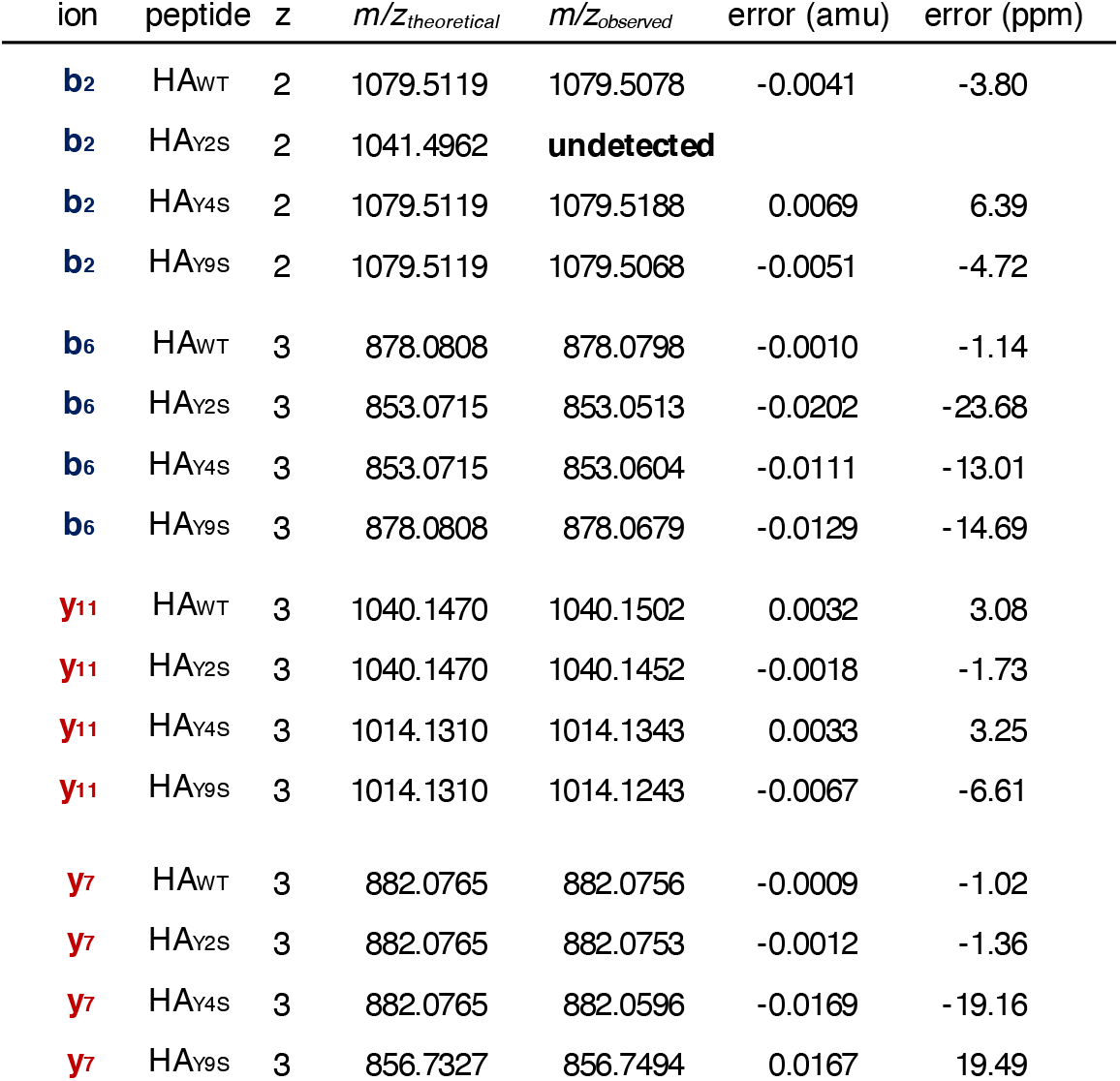
Mass Error Analysis for Ub_WT_ + HA Peptide Ser-Variants xl-fragment.

## References

1. Rowbotham, T. J. Preliminary report on the pathogenicity of Legionella pneumophila for freshwater and soil amoebae. J. Clin. Pathol. 33, 1179–1183 (1980).

2. Escoll, P., Rolando, M., Gomez-Valero, L. & Buchrieser, C. From amoeba to macrophages: exploring the molecular mechanisms of Legionella pneumophila infection in both hosts. Curr. Top. Microbiol. Immunol. 376, 1–34 (2013).

3. Swanson, M. S. & Isberg, R. R. Association of Legionella pneumophila with the macrophage endoplasmic reticulum. Infect. Immun. 63, 3609–3620 (1995).

4. Coers, J., Monahan, C. & Roy, C. R. Modulation of phagosome biogenesis by Legionella pneumophila creates an organelle permissive for intracellular growth. Nat. Cell Biol. 1, 451–453 (1999).

5. Glick, T. H. et al. Pontiac fever. An epidemic of unknown etiology in a health department: I. Clinical and epidemiologic aspects. Am. J. Epidemiol. 107, 149–160 (1978).

6. Berger, K. H. & Isberg, R. R. Two distinct defects in intracellular growth complemented by a single genetic locus in Legionella pneumophila. Mol. Microbiol. 7, 7–19 (1993).

7. Marra, A., Blander, S. J., Horwitz, M. A. & Shuman, H. A. Identification of a Legionella pneumophila locus required for intracellular multiplication in human macrophages. Proc. Natl. Acad. Sci. USA 89, 9607–9611 (1992).

8. Swanson, M. S. & Hammer, B. K. Legionella pneumophila pathogesesis: a fateful journey from amoebae to macrophages. Annu. Rev. Microbiol. 54, 567–613 (2000).

9. Isberg, R. R., O’Connor, T. J. & Heidtman, M. The Legionella pneumophila replication vacuole: making a cosy niche inside host cells. Nat. Rev. Microbiol. 7, 13–24 (2009).

10. Vincent, C. D. et al. Identification of the core transmembrane complex of the Legionella Dot/Icm type IV secretion system. Mol. Microbiol. 62, 1278–1291 (2006).

11. Laguna, R. K., Creasey, E. A., Li, Z., Valtz, N. & Isberg, R. R. A Legionella pneumophila-translocated substrate that is required for growth within macrophages and protection from host cell death. Proc. Natl. Acad. Sci. USA 103, 18745–18750 (2006).

12. Creasey, E. A. & Isberg, R. R. The protein SdhA maintains the integrity of the Legionella-containing vacuole. Proc. Natl. Acad. Sci. USA 109, 3481–3486 (2012).

13. Anand, I. S., Choi, W. Y. & Isberg, R. R. Components of the endocytic and recycling trafficking pathways interfere with the integrity of the Legionella-containing vacuole. Cell Microbiol. e13151 (2020). doi:10.1111/cmi.13151

14. Haenssler, E., Ramabhadran, V., Murphy, C. S., Heidtman, M. I. & Isberg, R. R. Endoplasmic Reticulum Tubule Protein Reticulon 4 Associates with the Legionella pneumophila Vacuole and with Translocated Substrate Ceg9. Infect. Immun. 83, 3479–3489 (2015).

15. Kotewicz, K. M. et al. A single legionella effector catalyzes a multistep ubiquitination pathway to rearrange tubular endoplasmic reticulum for replication. Cell Host Microbe 21, 169–181 (2017).

16. Hervet, E. et al. Protein kinase LegK2 is a type IV secretion system effector involved in endoplasmic reticulum recruitment and intracellular replication of Legionella pneumophila. Infect. Immun. 79, 1936–1950 (2011).

17. Luo, Z.-Q. & Isberg, R. R. Multiple substrates of the Legionella pneumophila Dot/Icm system identified by interbacterial protein transfer. Proc. Natl. Acad. Sci. USA 101, 841–846 (2004).

18. Ensminger, A. W. & Isberg, R. R. E3 ubiquitin ligase activity and targeting of BAT3 by multiple Legionella pneumophila translocated substrates. Infect. Immun. 78, 3905–3919 (2010).

19. Hsu, F. et al. The Legionella effector SidC defines a unique family of ubiquitin ligases important for bacterial phagosomal remodeling. Proc. Natl. Acad. Sci. USA 111, 10538–10543 (2014).

20. Belyi, Y., Tabakova, I., Stahl, M. & Aktories, K. Lgt: a family of cytotoxic glucosyltransferases produced by Legionella pneumophila. J. Bacteriol. 190, 3026–3035 (2008).

21. Shen, X. et al. Targeting eEF1A by a Legionella pneumophila effector leads to inhibition of protein synthesis and induction of host stress response. Cell Microbiol. 11, 911–926 (2009).

22. Barry, K. C., Fontana, M. F., Portman, J. L., Dugan, A. S. & Vance, R. E. IL-1α signaling initiates the inflammatory response to virulent Legionella pneumophila in vivo. J. Immunol. 190, 6329–6339 (2013).

23. Chambers, K. A. & Scheck, R. A. Bacterial virulence mediated by orthogonal post-translational modification. Nat. Chem. Biol. 16, 1043–1051 (2020).

24. O’Connor, T. J., Boyd, D., Dorer, M. S. & Isberg, R. R. Aggravating genetic interactions allow a solution to redundancy in a bacterial pathogen. Science 338, 1440–1444 (2012).

25. Blatt, S. P. et al. Nosocomial Legionnaires’ disease: aspiration as a primary mode of disease acquisition. Am. J. Med. 95, 16–22 (1993).

26. Bardill, J. P., Miller, J. L. & Vogel, J. P. IcmS-dependent translocation of SdeA into macrophages by the Legionella pneumophila type IV secretion system. Mol. Microbiol. 56, 90–103 (2005).

27. Shin, D. et al. Regulation of Phosphoribosyl-Linked Serine Ubiquitination by Deubiquitinases DupA and DupB. Mol. Cell 77, 164–179.e6 (2020).

28. Wan, M. et al. Deubiquitination of phosphoribosyl-ubiquitin conjugates by phosphodiesterase-domain-containing Legionella effectors. Proc. Natl. Acad. Sci. USA 116, 23518–23526 (2019).

29. Sulpizio, A. G., Minelli, M. E. & Mao, Y. Glutamylation of bacterial ubiquitin ligases by a legionella pseudokinase. Trends Microbiol. (2019). doi:10.1016/j.tim.2019.09.001

30. Gan, N. et al. Regulation of phosphoribosyl ubiquitination by a calmodulin-dependent glutamylase. Nature 572, 387–391 (2019).

31. Black, M. H. et al. Bacterial pseudokinase catalyzes protein polyglutamylation to inhibit the SidE-family ubiquitin ligases. Science 364, 787–792 (2019).

32. Bhogaraju, S. et al. Phosphoribosylation of ubiquitin promotes serine ubiquitination and impairs conventional ubiquitination. Cell 167, 1636–1649.e13 (2016).

33. Sheedlo, M. J. et al. Structural basis of substrate recognition by a bacterial deubiquitinase important for dynamics of phagosome ubiquitination. Proc. Natl. Acad. Sci. USA 112, 15090–15095 (2015).

34. Puvar, K. et al. Ubiquitin Chains Modified by the Bacterial Ligase SdeA Are Protected from Deubiquitinase Hydrolysis. Biochemistry 56, 4762–4766 (2017).

35. Qiu, J. et al. Ubiquitination independent of E1 and E2 enzymes by bacterial effectors. Nature 533, 120–124 (2016).

36. Kim, L. et al. Structural and Biochemical Study of the Mono-ADP-Ribosyltransferase Domain of SdeA, a Ubiquitylating/Deubiquitylating Enzyme from Legionella pneumophila. J. Mol. Biol. 430, 2843–2856 (2018).

37. Kalayil, S. et al. Insights into catalysis and function of phosphoribosyl-linked serine ubiquitination. Nature 557, 734–738 (2018).

38. Dong, Y. et al. Structural basis of ubiquitin modification by the Legionella effector SdeA. Nature 557, 674–678 (2018).

39. Akturk, A. et al. Mechanism of phosphoribosyl-ubiquitination mediated by a single Legionella effector. Nature 557, 729–733 (2018).

40. Hershko, A. Ciechanover A. The ubiquitin system. Annu Rev Biochem (1998).

41. Wang, Y. et al. Structural Insights into Non-canonical Ubiquitination Catalyzed by SidE. Cell 173, 1231–1243.e16 (2018).

42. Bhogaraju, S. et al. Inhibition of bacterial ubiquitin ligases by SidJ-calmodulin-catalysed glutamylation. Nature (2019). doi:10.1038/s41586-019-1440-8

43. Wong, K., Kozlov, G., Zhang, Y. & Gehring, K. Structure of the Legionella Effector, lpg1496, Suggests a Role in Nucleotide Metabolism. J. Biol. Chem. 290, 24727–24737 (2015).

44. Jeong, K. C., Sexton, J. A. & Vogel, J. P. Spatiotemporal regulation of a Legionella pneumophila T4SS substrate by the metaeffector SidJ. PLoS Pathog. 11, e1004695 (2015).

45. Machner, M. P. & Isberg, R. R. Targeting of host Rab GTPase function by the intravacuolar pathogen Legionella pneumophila. Dev. Cell 11, 47–56 (2006).

46. Yau, R. & Rape, M. The increasing complexity of the ubiquitin code. Nat. Cell Biol. 18, 579–586 (2016).

47. Behrends, C. & Harper, J. W. Constructing and decoding unconventional ubiquitin chains. Nat. Struct. Mol. Biol. 18, 520–528 (2011).

48. Kulathu, Y. & Komander, D. Atypical ubiquitylation - the unexplored world of polyubiquitin beyond Lys48 and Lys63 linkages. Nat. Rev. Mol. Cell Biol. 13, 508–523 (2012).

