## Supplementary Information for "Members of the *Legionella pneumophila* Sde Family Target Tyrosine Residues for Phosphoribosyl-Linked Ubiquitination"

### Electronic Supplementary Information

**Materials and Methods**

General Materials 2
Protein Purification and Peptide Synthesis 2
Rtn4 Ubiquitination Using Two-step SdeC Bead Assay 3
HA-Ub Polymerization Assay and HA Peptide Ubiquitination Assays 3
Polyacrylamide Gel Electrophoresis and Western Blotting 3
FPLC Purification and Size Exclusion Chromatography 3
General Protocol for Trypsin Digestion of ADP-ribosylated Ub and Crosslinked Proteins 3
Mass Spectrometry 3

**Supplementary Figures and Tables**

**Figure S1** HA-Ub polymerization is dependent on HA tag 5
**Table S1** Mass error anlysis for HA-Ub xl-fragment 6
**Table S2** Mass Error Analysis for $Ub_{WT}$ + HA Peptide Phe-Variants xl-fragment. 7
**Table S3** Mass Error Analysis for $Ub_{WT}$ + $HAY_{249S}$ xl-fragment 8
**Figure S2** SdeC does not have an intrinsic preference for Ser over Tyr linkages 9
**Table S4** Mass Error Analysis for $Ub_{WT}$ + HA Peptide Ser-Variants xl-fragment 10
**Table S5** Plasmids used in this study 11
**Table S6** Peptides used in this study 11

**Supplementary References** 12

**General Materials.** All chemical reagents and solvents were of analytical grade, obtained from commercial suppliers and used without further purification unless otherwise noted.

**Protein Purification and Peptide Synthesis.** SdeC variants were purified as described previously.[1,2] *Escherichia coli* strain BL21 (DE3) strain carrying bacterial expression vector pQE-80L (N-terminal Poly His tag) with either full length SdeC WT or point mutants, each containing a C-terminal StrepII tag, was used for protein expression. Bacteria cultures were grown in 2X Yeast Tryptone broth (2xYT) overnight with 100 μg/ml ampicillin, back diluted and grown until A600 = 0.6, at which point 1mM IPTG (Sigma) was added for 4-6 hours to induce protein expression. Cells were collected by centrifugation, resuspended in 2% of the culture volume with lysis buffer (50mM $NaH_2PO_4$, 300 mM NaCl, 10 mM imidazole, pH 7.6), then subjected to 4-5 freeze-thaw cycles before sonication. Soluble proteins were purified from collected supernatants using Ni-NTA agarose resin (Thermo Fisher) followed by Strep-tactin resin (IBA) affinity chromatography according to manufacturer's protocol. Purified proteins were dialyzed into 20 mM Tris, 10 mM NaCl, pH 7.4 (1X ART buffer), and concentrated with Amicon Ultra Centrifuge filters (30K MWCO). Protein concentrations were determined by Bradford assay (Bio-Rad) using BSA as standard. Concentrated proteins were then mixed with 10% glycerol, aliquoted, and stored at -80°C. Purified GST-Rtn4 was purchased from MRCPPU. Human recombinant monomeric Ub and HA-Ub were purchased from Boston Biochem (R&D systems). All HA peptide variants were synthesized in Tufts University Core Facility. All plasmids and peptides in this study are further described in **Tables S5 & S6**.

**Rtn4 Ubiquitination Using Two-step SdeC Bead Assay.** *In vitro* Rtn4 ubiquitination assays were described previously.[1] For linking recombinant SdeC derivatives to agarose beads, Ni-NTA agarose resin (Thermo Fisher) was washed 3X with ART buffer and mixed with $His_6$-SdeC followed by incubation for 1 h at 4 °C. Then, the SdeC-bound agarose resin was washed 3X with 1X ART buffer before proceeding to the two-step bead assay. During preincubation, SdeC-bound agarose beads were added to 10 mM Ub and 100 mM ε-NAD in 1X ART buffer (SdeC final concentration 50 nM), and incubated for 1 h at 37 °C. After incubation, the beads were removed from reaction by spin filter columns (Pierce spin cups, paper filter that was pre-wet with 1X ART buffer). The supernatant containing modified Ub and excess NAD were mixed with 400 nM GST-HA-Rtn4 and fresh unbound SdeC variants, as indicated, then incubated for another hour at 37 °C. Reactions were terminated by addition of reducing loading buffer and boiling.

**HA-Ub Polymerization Assay and HA Peptide Ubiquitination Assays.** For HA-Ub polymerization assays, 1 μM HA-Ub was mixed with 100 μM Nicotinamide 1, N6-ethenoadenine dinucleotide (ε-NAD, Sigma) or β-Nicotinamide adenine dinucleotide (β-NAD, Santa Cruz Biotech/Sigma), and recombinant SdeC at final concentration of 20 nM. The reaction mix was incubated at 37 °C in 1X ART buffer for 2 h. For HA peptide ubiquitination assays, 12.5 μM HA derivative peptides were mixed with 10 μM monomeric Ub, and incubated with 20 nM SdeC and 250 μM β-NAD at 37°C in 1X ART buffer for 1 h. All reactions for gel analysis were terminated by either addition of reducing loading buffer and boiling, or flash freezing in liquid $N_2$ for storage prior to further analysis. For samples analyzed by mass spectrometry (MS), excess salts and NAD were removed via buffer exchange, using either Illustra NAP-5 Columns (Cytiva Life sciences) or Amicon Ultra Centrifuge filters, 3K MWCO (MilliporeSigma), eluting in water.

**Polyacrylamide Gel Electrophoresis and Western Blotting.** After assays for HA-Ub crosslinking or pR-Ub modification, samples boiled in reducing SDS loading buffer were fractionated by sodium dodecyl sulfate polyacrylamide gel electrophoresis (SDS-PAGE). Assays using Ub as a substrate, as well as HA-Ub and HA-peptides, were fractionated on 15% gels, while the GST-Rtn4 reactions were fractionated on 5% gels followed by either Coomassie Brilliant Blue staining or silver staining. Silver stains were performed as described in manufacturer's protocol (Invitrogen SilverQuest Silver Staining kit). The mouse monoclonal anti-Ub antibody (mAb P4D1, 1:500) and anti-ethenoadenosine mAb (1G4; 1:500) were purchased from Santa Cruz Biotechnology. Anti-mouse IgG (H+L) (DyLight™ 800 4X PEG Conjugate) was purchased from Cell Signaling Technology (1:20,000). LI-COR Odyssey CLx was used for Western blot signal detection.

Analysis was completed using multiple programs. Agilent MassHunter Bioconfirm (v. B.07.00) was used for generation of intact protein deconvolutions. Bioconfirm was also used to generate extracted ion chromatograms (EIC) and additional mass spectra. Exact masses were calculated using ChemDraw Prime (PerkinElmer, v. 16.0). ChemDraw also enabled identification of unique fragmentation patterns. Posttranslational modifications (PTMs) were identified using PEAKS Studio (v. 7.5) software. Utilizing PEAKS Search feature, preliminary analysis enabled 0.2 Da error tolerance for a tryptic digestion with non-specific cleavage allowed at either end of the peptides and up to 3 missed cleavages. Unspecified PTMs and common mutations were searched to match Ubiquitin or pR-HA-Ub sequences. For verification, new PTMs were created in the software and analysis repeated with greater stringencies to confirm MS/MS matching[3,4]. Mass error analysis was completed by matching observed fragment ions with predicted fragmentations generated in ChemDraw. Error is reported in ppm as well as amu. Figures created in Adobe Illustrator.

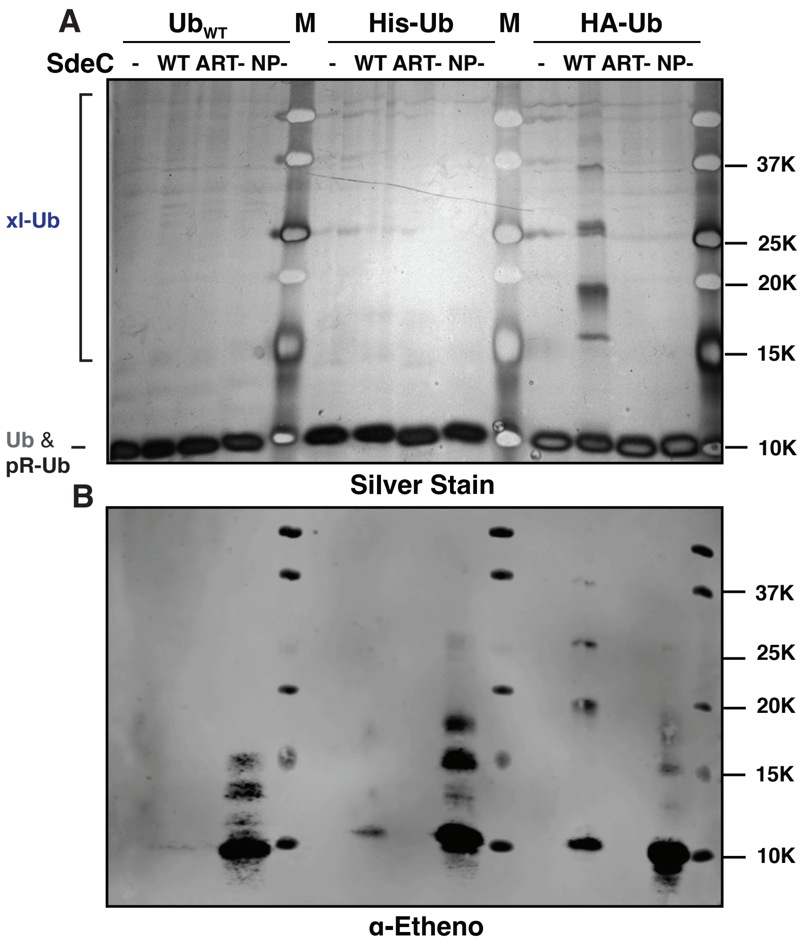


**Figure S1. HA-Ub polymerization is dependent on HA tag. Linked to Fig. 2 A)** WT Ub, His-Ub, HA-Ub were incubated with $NAD^+$ and SdeC at 37°C for 2 h, then fractionated on a SDS-PAGE gel, followed by silver staining. **B)** WT Ub, His-Ub, HA-Ub were incubated with ε-$NAD^+$ in presence of purified SdeC variants at 37C for 2 hr, then fractionated on a 15% SDS-PAGE gel, followed by silver staining (top) or anti-ethenoadenosine staining (bottom).

**Table S1. Mass Error Anlysis for HA-Ub xl-fragment.**
Mass error analysis for identified and assigned fragment ions labeled in Main Text **Figure 3**.

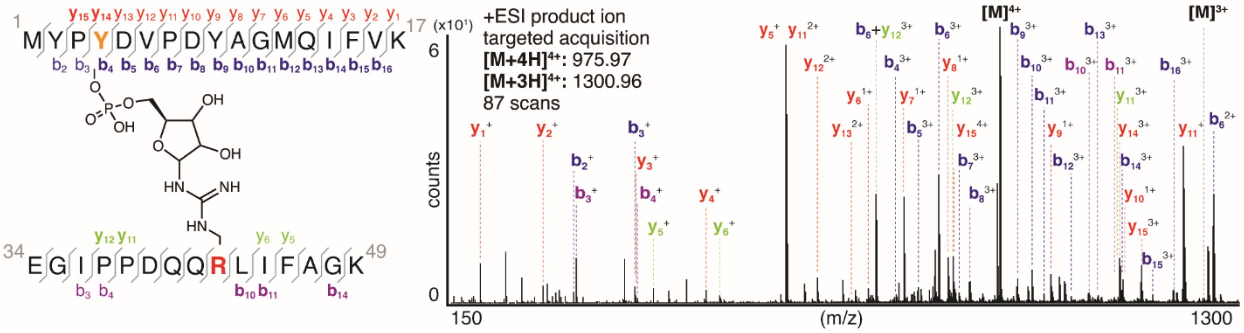


**HA tag (1-17) fragmentations:**

| ion | z | $m/z_{theoretical}$ | $m/z_{observed}$ | error (amu) | error (ppm) |
|---|---|---|---|---|---|
| $b_{2}$ | 1 | 296.1090 | 296.1076 | -0.0014 | -4.73 |
| $b_{3}$ | 1 | 393.1717 | 393.1734 | 0.0017 | 4.32 |
| $b_{4}$ | 2 | 1210.0738 | 1210.0714 | -0.0024 | -1.98 |
| $b_{4}$ | 3 | 807.3846 | 807.3748 | -0.0098 | -12.14 |
| $b_{5}$ | 2 | 1267.5873 | 1267.5795 | -0.0078 | -6.15 |
| $b_{5}$ | 3 | 845.3940 | 845.3875 | -0.0065 | -7.69 |
| $b_{6}$ | 2 | 1317.6176 | 1317.6194 | 0.0018 | 1.37 |
| $b_{6}$ | 3 | 878.0808 | 878.0810 | 0.0002 | 0.23 |
| $b_{7}$ | 3 | 910.7662 | 910.7623 | -0.0039 | -4.28 |
| $b_{8}$ | 3 | 949.1085 | 949.1054 | -0.0031 | -3.27 |
| $b_{9}$ | 3 | 1003.4645 | 1003.4618 | -0.0027 | -2.69 |
| $b_{10}$ | 3 | 1027.1435 | 1027.1440 | 0.0005 | 0.49 |
| $b_{11}$ | 3 | 1046.1506 | 1046.1446 | -0.0060 | -5.74 |
| $b_{12}$ | 3 | 1090.1653 | 1090.1652 | -0.0001 | -0.09 |
| $b_{13}$ | 3 | 1132.8514 | 1132.8465 | -0.0049 | -4.33 |
| $b_{14}$ | 3 | 1170.2117 | 1170.2082 | -0.0035 | -2.99 |
| $b_{15}$ | 3 | 1219.2345 | 1219.2450 | 0.0105 | 8.61 |
| $b_{16}$ | 3 | 1252.5917 | 1252.5903 | -0.0014 | -1.12 |

| ion | z | $m/z_{theoretical}$ | $m/z_{observed}$ | error (amu) | error (ppm) |
|---|---|---|---|---|---|
| $y_{15}$ | 3 | 1201.9167 | 1201.9062 | -0.0105 | -8.74 |
| $y_{15}$ | 4 | 901.6893 | 901.6833 | -0.0060 | -6.65 |
| $y_{14}$ | 3 | 1169.8981 | 1169.8777 | -0.0204 | -17.44 |
| $y_{13}$ | 1 | 1482.7253 | 1482.7178 | -0.0075 | -5.06 |
| $y_{13}$ | 2 | 741.3624 | 741.3785 | 0.0161 | 21.72 |
| $y_{12}$ | 1 | 1367.6984 | 1367.6971 | -0.0013 | -0.95 |
| $y_{12}$ | 2 | 684.3506 | 684.3550 | 0.0044 | 6.43 |
| $y_{11}$ | 1 | 1269.6333 | 1269.6364 | 0.0031 | 2.44 |
| $y_{11}$ | 2 | 635.3232 | 635.3203 | -0.0029 | -4.56 |
| $y_{10}$ | 1 | 1172.5805 | 1172.5830 | 0.0025 | 2.13 |
| $y_{9}$ | 1 | 1057.5536 | 1057.5511 | -0.0025 | -2.36 |
| $y_{8}$ | 1 | 893.4869 | 893.4896 | 0.0027 | 3.02 |
| $y_{7}$ | 1 | 822.4498 | 822.4543 | 0.0045 | 5.47 |
| $y_{6}$ | 1 | 765.4283 | 765.4322 | 0.0039 | 5.10 |
| $y_{5}$ | 1 | 634.3879 | 634.3888 | 0.0009 | 1.42 |
| $y_{4}$ | 1 | 506.3293 | 506.3335 | 0.0042 | 8.29 |
| $y_{3}$ | 1 | 393.2452 | 393.2486 | 0.0034 | 8.65 |
| $y_{2}$ | 1 | 246.1768 | 246.1805 | 0.0037 | 15.03 |
| $y_{1}$ | 1 | 147.1084 | 147.1108 | 0.0024 | 16.31 |

**R42 Tryptic product (34-49) fragmentations:**

| ion | z | $m/z_{theoretical}$ | $m/z_{observed}$ | error (amu) | error (ppm) |
|---|---|---|---|---|---|
| $b_{3}$ | 1 | 301.1633 | 301.1610 | -0.0023 | -7.64 |
| $b_{4}$ | 1 | 398.2160 | 398.2048 | -0.0112 | -28.13 |
| $b_{10}$ | 3 | 1122.5271 | 1122.5177 | -0.0094 | -8.37 |
| $b_{11}$ | 3 | 1160.5468 | 1160.5504 | 0.0036 | 3.10 |
| $b_{14}$ | 3 | 1252.5917 | 1252.5903 | -0.0014 | -1.12 |

| ion | z | $m/z_{theoretical}$ | $m/z_{observed}$ | error (amu) | error (ppm) |
|---|---|---|---|---|---|
| $y_{12}$ | 4 | 901.1808 | 901.1825 | 0.0017 | 1.89 |
| $y_{11}$ | 3 | 1169.2222 | 1169.2103 | -0.0119 | -10.18 |
| $y_{6}$ | 1 | 535.3194 | 535.3090 | -0.0104 | -19.43 |
| $y_{5}$ | 1 | 422.2354 | 422.2308 | -0.0046 | -10.89 |

**Double N-terminal Pro fragmentation:**

| ion | z | $m/z_{theoretical}$ | $m/z_{observed}$ | error (amu) | error (ppm) |
|---|---|---|---|---|---|
| $b_{6}+y_{12}$ | 3 | 778.3633 | 778.3646 | 0.0013 | 1.67 |

**Table S2: Mass Error Analysis for Ub$_{WT}$ + HA Peptide Phe-Variants xl-fragment**
Mass error analysis for identified and assigned fragment ions labeled in **Figure 5**.

| ion | peptide | z | $m/z_{theoretical}$ | $m/z_{observed}$ | error (amu) | error (ppm) |
|---|---|---|---|---|---|---|
| $b_2$ | $HA_{WT}$ | 2 | 1079.5119 | 1079.508 | -0.0041 | -3.80 |
| $b_2$ | $HA_{Y2F}$ | 2 | 1071.5144 | **undetected** | | |
| $b_2$ | $HA_{Y4F}$ | 2 | 1079.5119 | 1079.5103 | -0.0016 | -1.48 |
| $b_2$ | $HA_{Y9F}$ | 2 | 1079.5119 | 1079.4992 | -0.0127 | -11.76 |
| $y_{11}$ | $HA_{WT}$ | 3 | 1040.1470 | 1040.1502 | 0.0032 | 3.08 |
| $y_{11}$ | $HA_{Y2F}$ | 3 | 1040.1470 | 1040.1480 | 0.0010 | 0.96 |
| $y_{11}$ | $HA_{Y4F}$ | 3 | 1034.8154 | 1034.8160 | 0.0006 | 0.58 |
| $y_{11}$ | $HA_{Y9F}$ | 3 | 1034.8154 | 1034.8190 | 0.0036 | 3.48 |
| $b_6$ | $HA_{WT}$ | 3 | 878.0808 | 878.0798 | -0.0010 | -1.14 |
| $b_6$ | $HA_{Y2F}$ | 3 | 873.0836 | 873.0877 | 0.0041 | 4.70 |
| $b_6$ | $HA_{Y4F}$ | 3 | 873.0836 | 873.0908 | 0.0072 | 8.25 |
| $b_6$ | $HA_{Y9F}$ | 3 | 878.0808 | 878.0822 | 0.0014 | 1.59 |
| $y_7$ | $HA_{WT}$ | 3 | 882.0765 | 882.0756 | -0.0009 | -1.02 |
| $y_7$ | $HA_{Y2F}$ | 3 | 882.0765 | 882.0752 | -0.0013 | -1.47 |
| $y_7$ | $HA_{Y4F}$ | 3 | 882.0765 | 882.0752 | -0.0013 | -1.47 |
| $y_7$ | $HA_{Y9F}$ | 3 | 877.0808 | **undetected** | | |

**Table S3: Mass Error Analysis for $Ub_{WT}$ + $HAY_{249S}$ xl-fragment.**

Mass error analysis for identified and assigned fragment ions labeled in **Main Text Figure 6**.

| ion | z | $m/z_{theoretical}$ | $m/z_{observed}$ | error (amu) | error (ppm) |
|---|---|---|---|---|---|
| $b_2$ | 1 | 221.0935 | 221.0910 | -0.0025 | -11.31 |
| $b_3$ | 1 | 317.1404 | 317.1359 | -0.0045 | -14.19 |
| $b_4$ | 1 | 405.1758 | 405.1754 | -0.0004 | -0.99 |
| $b_5$ | 1 | 519.1994 | 519.1908 | -0.0086 | -16.56 |
| $b_5$ | 3 | 618.2678 | 618.2570 | -0.0108 | -17.47 |
| $b_{10}$ | 3 | 951.1122 | 951.1153 | 0.0031 | 3.26 |
| $b_{11}$ | 3 | 970.1193 | 970.1133 | -0.0060 | -6.18 |
| $b_{12}$ | 3 | 1014.4684 | 1014.4676 | -0.0008 | -0.79 |
| $y_{12}$ | 3 | 1018.4690 | 1018.4547 | -0.0143 | -14.04 |
| $y_{11}$ | 3 | 989.4595 | 989.4663 | 0.0068 | 6.87 |
| $y_{10}$ | 3 | 956.4396 | 956.4321 | -0.0075 | -7.84 |
| $y_9$ | 3 | 927.4290 | 927.4353 | 0.0063 | 6.79 |
| $y_8$ | 3 | 889.7555 | 889.7499 | -0.0056 | -6.29 |
| $y_7$ | 3 | 857.0672 | 857.0643 | -0.0029 | -3.38 |
| $y_3$ | 1 | 334.1266 | 334.1282 | 0.0016 | 4.79 |
| $y_2$ | 1 | 276.1018 | 276.1076 | 0.0058 | 21.01 |

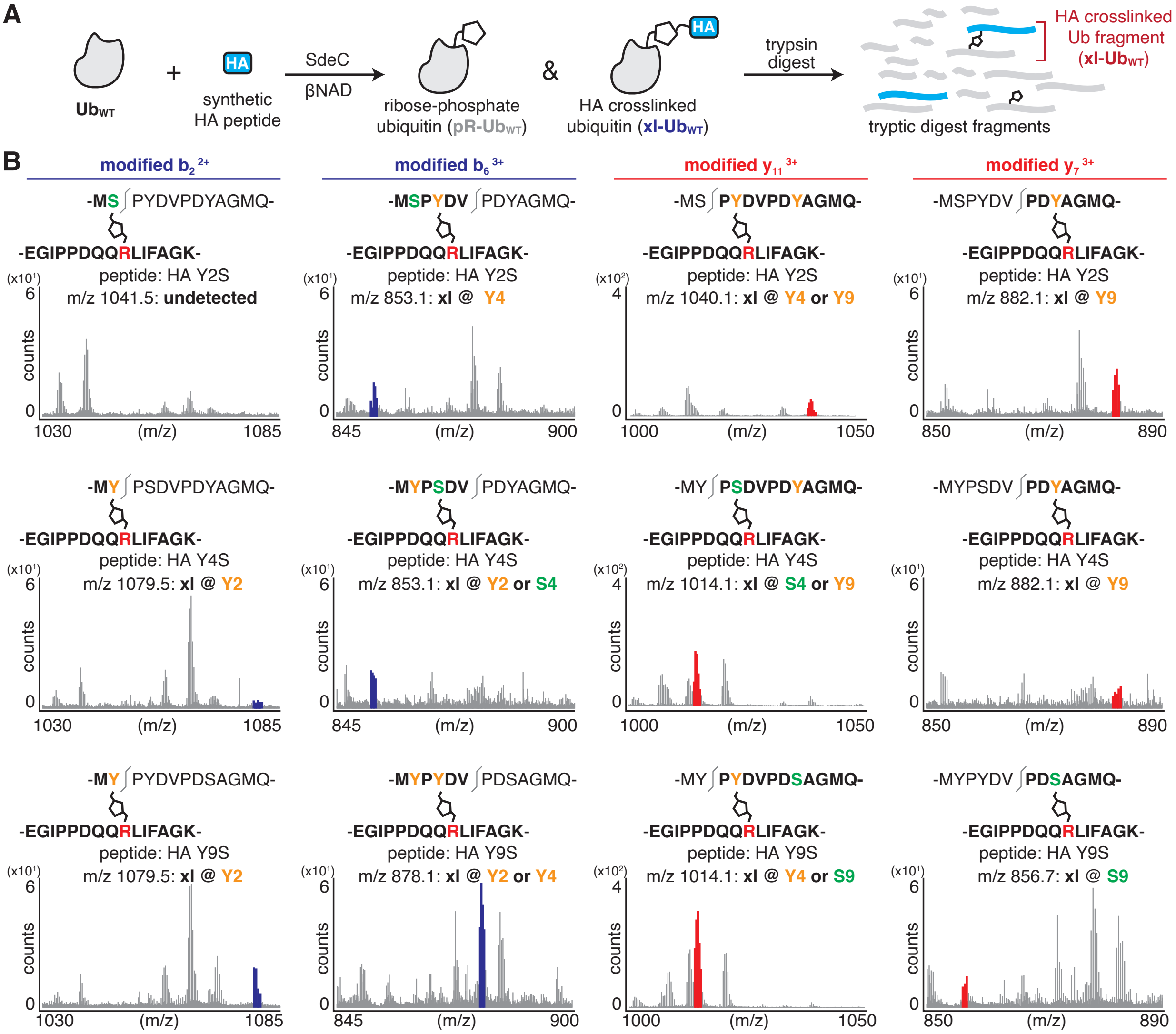


**Figure S2. SdeC does not have an intrinsic preference for Ser over Tyr linkages. A)** Scheme depicting the SdeC-catalyzed phosphoribosylation of ubiquitin (wtUb) and its crosslinking to synthetic HA peptides in the presence of βNAD. **B)** After tryptic digest, peptide mixtures were analyzed using LC-MS/MS. Key fragmentation ions are highlighted, indicating that crosslinking can occur at multiple positions within the synthetic HA peptide. The m/z for the modified $b_2$ ion, which would suggest crosslinking at Y2 or S2, is absent in HA$_{Y2S}$, but present in HA$_{Y4S}$ and HA$_{Y9S}$. The modified $b_6$ ion is present in all three peptide variants, and indicates crosslinking either at Y4 for HA$_{Y2S}$. For HA$_{Y4S}$ , the presence of this ion suggests that the site of modification may be at either Y2 or S4. For HA$_{Y4S}$, the presence of this ion suggests that the site of modification may be at either Y2 or Y4. Similarly, a modified $y_{11}$ ion is present in all peptide variants, though this ion would suggest two possible sites of crosslinking, each highlighted in orange, as indicated for each spectra. Finally, the modified $y_7$ ion indicates crosslinking at Y9 or S9 only. We found that the only definitive evidence of modification at serine for these peptides occurred at S9 for HA$_{Y9S}$. For HA$_{Y2S}$ and HA$_{Y4S}$, this ion is present and indicates that crosslinking occurs at Y9.

**Table S4: Mass Error Analysis for $Ub_{WT}$ + HA Peptide Ser-Variants xl-fragment.** Mass error analysis for identified and assigned fragment ions labeled in **Figure S2**.

| ion | peptide | z | $m/z_{theoretical}$ | $m/z_{observed}$ | error (amu) | error (ppm) |
|---|---|---|---|---|---|---|
| $b_2$ | $HA_{WT}$ | 2 | 1079.5119 | 1079.5078 | -0.0041 | -3.80 |
| $b_2$ | $HA_{Y2S}$ | 2 | 1041.4962 | **undetected** | | |
| $b_2$ | $HA_{Y4S}$ | 2 | 1079.5119 | 1079.5188 | 0.0069 | 6.39 |
| $b_2$ | $HA_{Y9S}$ | 2 | 1079.5119 | 1079.5068 | -0.0051 | -4.72 |
| $b_6$ | $HA_{WT}$ | 3 | 878.0808 | 878.0798 | -0.0010 | -1.14 |
| $b_6$ | $HA_{Y2S}$ | 3 | 853.0715 | 853.0513 | -0.0202 | -23.68 |
| $b_6$ | $HA_{Y4S}$ | 3 | 853.0715 | 853.0604 | -0.0111 | -13.01 |
| $b_6$ | $HA_{Y9S}$ | 3 | 878.0808 | 878.0679 | -0.0129 | -14.69 |
| $y_{11}$ | $HA_{WT}$ | 3 | 1040.1470 | 1040.1502 | 0.0032 | 3.08 |
| $y_{11}$ | $HA_{Y2S}$ | 3 | 1040.1470 | 1040.1452 | -0.0018 | -1.73 |
| $y_{11}$ | $HA_{Y4S}$ | 3 | 1014.1310 | 1014.1343 | 0.0033 | 3.25 |
| $y_{11}$ | $HA_{Y9S}$ | 3 | 1014.1310 | 1014.1243 | -0.0067 | -6.61 |
| $y_7$ | $HA_{WT}$ | 3 | 882.0765 | 882.0756 | -0.0009 | -1.02 |
| $y_7$ | $HA_{Y2S}$ | 3 | 882.0765 | 882.0753 | -0.0012 | -1.36 |
| $y_7$ | $HA_{Y4S}$ | 3 | 882.0765 | 882.0596 | -0.0169 | -19.16 |
| $y_7$ | $HA_{Y9S}$ | 3 | 856.7327 | 856.7494 | 0.0167 | 19.49 |

**Table S5.** Plasmids used in this study.

| **Plasmids** | | | |
|---|---|---|---|
| **No.** | **Plasmid Name** | **Details** | **Protein activity** |
| **1** | pQE-80L-SdeC$_{WT}$[1] | pQE-80L *sdeC*, N-terminus 6XHis tag, C-terminus StrepII tag | SdeC WT |
| **2** | pQE-80L-SdeC$_{H416A}$[1] | pQE-80L *sdeC* H416A, N-terminus 6XHis tag, C-terminus StrepII tag | SdeC NP- |
| **3** | pQE-80L-SdeC$_{E859A}$[1] | pQE-80L *sdeC* E859A, N-terminus 6XHis tag, C-terminus StrepII tag | SdeC ART- |

**Table S6.** Peptides used in this study.

| **Peptides** | | | |
|---|---|---|---|
| **No.** | Peptide Name | Sequence | Exact Mass (mono) |
| **1** | HA$_{WT}$ | MYPYDVPDYAGMQ-$NH_2$ | 1547.631 |
| **2** | HA$_{Y2F}$ | M**F**PYDVPDYAGMQ-$NH_2$ | 1531.637 |
| **3** | HA$_{Y4F}$ | MYP**F**DVPDYAGMQ-$NH_2$ | 1531.637 |
| **4** | HA$_{Y9F}$ | MYPYDVPD**F**AGMQ-$NH_2$ | 1531.637 |
| **5** | HA$_{Y249F}$ | M**F**P**F**DVPD**F**AGMQ-$NH_2$ | 1499.647 |
| **6** | HA$_{Y2S}$ | M**S**PYDVPDYAGMQ-$NH_2$ | 1471.600 |
| **7** | HA$_{Y4S}$ | MYP**S**DVPDYAGMQ-$NH_2$ | 1471.600 |
| **8** | HA$_{Y9S}$ | MYPYDVPD**S**AGMQ-$NH_2$ | 1471.600 |
| **9** | HS$_{Y249S}$ | M**S**P**S**DVPD**S**AGMQ-$NH_2$ | 1319.538 |

**Supplementary References**


1. Kotewicz, K. M. *et al.* A single legionella effector catalyzes a multistep ubiquitination pathway to rearrange tubular endoplasmic reticulum for replication. *Cell Host Microbe* **21,** 169–181 (2017).
2. Machner, M. P. & Isberg, R. R. Targeting of host Rab GTPase function by the intravacuolar pathogen Legionella pneumophila. *Dev. Cell* **11,** 47–56 (2006).
3. Khatun, J., Ramkissoon, K. & Giddings, M. C. Fragmentation characteristics of collision-induced dissociation in MALDI TOF/TOF mass spectrometry. *Anal. Chem.* **79,** 3032–3040 (2007).
4. Brenton, A. G. & Godfrey, A. R. Accurate mass measurement: terminology and treatment of data. *J Am Soc Mass Spectrom* **21,** 1821–1835 (2010).
